# PHIStruct: Improving phage-host interaction prediction at low sequence similarity settings using structure-aware protein embeddings

**DOI:** 10.1101/2024.08.24.609479

**Authors:** Mark Edward M. Gonzales, Jennifer C. Ureta, Anish M.S. Shrestha

## Abstract

**Motivation:** Recent computational approaches for predicting phage-host interaction have explored the use of sequence-only protein language models to produce embeddings of phage proteins without manual feature engineering. However, these embeddings do not directly capture protein structure information and structure-informed signals related to host specificity.

**Result:** We present PHIStruct, a multilayer perceptron that takes in structure-aware embeddings of receptor-binding proteins, generated via the structure-aware protein language model SaProt, and then predicts the host from among the ESKAPEE genera. Compared against recent tools, PHIStruct exhibits the best balance of precision and recall, with the highest and most stable F1 score across a wide range of confidence thresholds and sequence similarity settings. The margin in performance is most pronounced when the sequence similarity between the training and test sets drops below 40%, wherein, at a relatively high-confidence threshold of above 50%, PHIStruct presents a 7% to 9% increase in class-averaged F1 over machine learning tools that do not directly incorporate structure information, as well as a 5% to 6% increase over BLASTp.

**Availability and Implementation:** The data and source code for our experiments and analyses are available at https://github.com/bioinfodlsu/PHIStruct.

## 1 Introduction

Described as a silent pandemic, antimicrobial resistance is among the foremost threats to public health. If left unaddressed, it is estimated that, by 2050, the number of deaths linked to antimicrobial resistance will reach 10 million annually and the global economic cost will exceed $100 trillion [63, 56]. As alternatives to conventional antibiotics, phages have recently been used in phage therapy to treat bacterial infections [29, 24] and in phage steering to re-sensitize pathogens to antibiotics [7]. The bactericidal action of phages has also been leveraged in various settings outside the clinical domain, including crop protection [53], oilfield reservoir souring prevention [45], and corrosion mitigation [25].

Identifying phage-host pairs is crucial in actualizing these applications; however, *in vitro* experiments can be time-consuming and expensive. In order to accelerate the shortlisting of candidate pairs, several *in silico* methods that capitalize on alignment-based and alignment-free techniques have been developed [31, 44]. Alignment-based approaches [69, 18, 49] focus on finding shared genomic regions between phages and putative hosts, whereas alignment-free approaches [12, 38, 37, 68, 41, 39] hinge on features such as sequence composition and co-abundance profiles of phages and their hosts.

In our previous work [23], we explored the use of representation learning to convert receptor-binding proteins (RBPs) into dense vector representations (embeddings) without the need for manual feature engineering. Located at the distal end of phages, RBPs, such as tailspike proteins, adsorb to receptors on the surface of the host bacteria and are thus key determinants of phage-host specificity [35, 20]. To generate the embeddings, we explored different protein language models, such as ESM-1b [47], ProtT5 [21], and SeqVec [27]. These models were pretrained in a self-supervised fashion on sequences from large-scale protein databases [6, 55, 50, 52] and have been shown to capture relevant physicochemical characteristics [21, 47, 27, 40]. We found that this approach presents performance improvements over using manually feature-engineered sequence properties.

From a biological perspective, additional benefit may be gained by incorporating structure information, as underscored by its role in determining function and elucidating protein-protein interaction signatures, such as complementary binding surfaces and molecular forces [46, 42]. In the particular context of phage-host interaction, recent studies have investigated the structure of RBPs to obtain insights into the mechanisms underlying host recognition and binding machinery. The structural diversity of these proteins has been found to reflect the evolutionary pressure to adapt to the surface proteins of bacteria occupying the same ecological niche [26, 15, 19].

Indeed, we found that phages infecting the same host genus have RBPs that are structurally more similar than those infecting different host genera, particularly at low sequence similarity settings. To see this, we paired RBPs targeting hosts from the ESKAPEE genera, which are among the leading causes of nosocomial infections worldwide [62]. The pairing was done such that the sequence similarity between the RBPs in each pair is below 40%. Using US-align [67], we then computed the root mean square deviation (RMSD) between their ColabFold [43]-predicted structures. Afterwards, for every genus, we constructed two groups, each with 500 randomly sampled RBP pairs. In the first group, both RBPs in each pair target the same genus of interest. In the second group, one of the RBPs in each pair targets the genus of interest, while the other RBP targets a different genus. Performing a Mann-Whitney U test showed that, for all but one genus, the distribution of the RMSD scores between these two groups is statistically significant at a *p*-value cutoff of 0.05 (Table A6).

These point towards the utility of integrating structure information in predicting phage-host interaction. A limitation of sequence-only protein language models (i.e., those pretrained only on sequences, without explicitly involving structure data) is that they do not directly capture interactions between residues that are in close proximity when the three-dimensional conformation is considered. This has motivated the development of structure-aware protein language models such as ProstT5 [28], SaProt [54], and PST [16], which adopt a custom vocabulary for encoding structure information [28, 54] or augment the architecture of an existing sequence-only model in order to inject structure bias into the output embeddings [16]. While structure-aware embeddings have been used for downstream benchmark tasks such as thermostability, metal ion binding, and protein function prediction [28, 54, 16], their application to computationally predicting phage-host pairs remains unexplored.

In this paper, we introduce PHIStruct, a deep learning model for phage-host interaction prediction that uses structure-aware embeddings of receptor-binding proteins for phage-host interaction prediction. We focused our scope on hosts belonging to the ESKAPEE genera (*Enterococcus*, *Staphylococcus*, *Klebsiella*, *Acinetobacter*, *Pseudomonas*, *Enterobacter*, and *Escherichia*), which include bacteria that are known to exhibit multidrug resistance [8, 36] and are among the priority pathogens identified by the World Health Organization [64]. Our contributions are as follows:

- We constructed a dataset of protein structures, computationally predicted via ColabFold [43], of 7,627 non-redundant (i.e., with duplicates removed) receptor-binding proteins from 3,350 phages that target hosts from the ESKAPEE genera. We identified these receptor-binding proteins based on GenBank annotations. For phage sequences without GenBank annotations, we employed a pipeline that uses the viral protein library PHROG [57] and the machine learning model PhageRBPdetect [13]. We also applied this pipeline to phages infecting other host genera, expanding our dataset to 19,081 nonredundant receptor-binding proteins from 8,525 phages across 238 host genera.
- We fed the ColabFold-predicted structures to the structure-aware protein language model SaProt [54] in order to generate embeddings of the receptor-binding proteins. We then trained a two-hidden-layer perceptron that takes in these structure-aware embeddings as input and predicts the host genus.
- We found that our model, PHIStruct, presents improvements over state-of-the-art tools that take in sequence-only protein embeddings and feature-engineered genomic and protein sequence properties, as well as BLASTp — especially as the sequence similarity between the training and test set entries decreases. Further evaluation highlights its ability to make high-confidence predictions without heavily compromising between precision and recall. When the sequence similarity drops below 40% and the confidence threshold is set to above 50%, our tool outperforms machine learning tools that do not directly integrate structure information by 7% to 9% and BLASTp by 5% to 6% in terms of class-averaged F1.

## 2 Methods

An overview of our methodology is provided in Figure 1. The steps are described in detail in the subsequent subsections.

**Figure 1:**
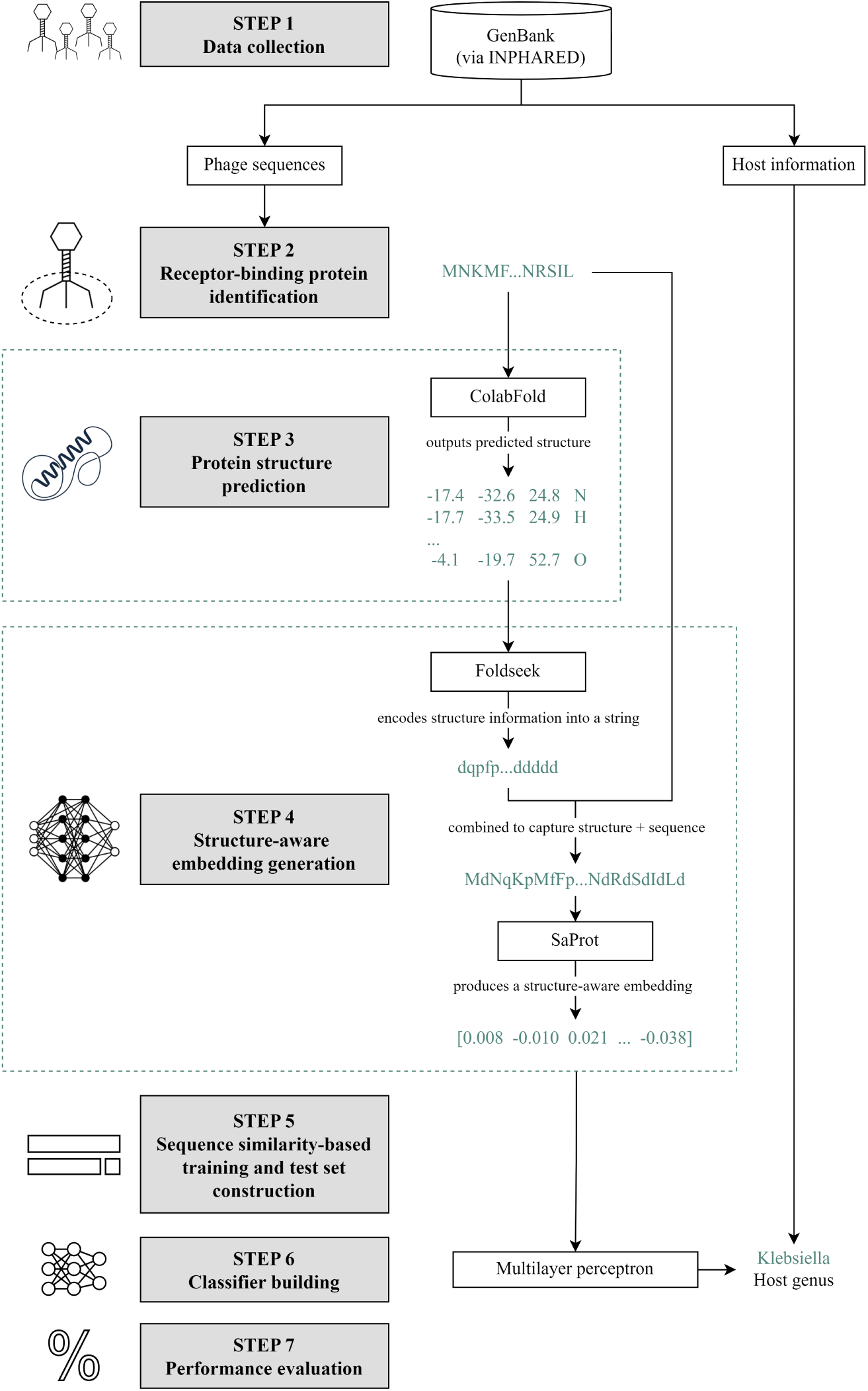
Methodology. Step 1: We collected phage genome and protein sequences from GenBank [11] using INPHARED [17]. Step 2: Receptor-binding proteins (RBPs) were identified following the methodology in our previous work [23]. Step 3: We fed the RBP sequences to ColabFold [43] to predict their structures. Step 4: The proteins, alongside their predicted structures, were fed to SaProt. For a protein of length *n*, the input to SaProt is ⟨(*r*_1_*, f*_1_), (*r*_2_*, f*_2_), (*r*_3_*, f*_3_) *, …,* (*r_n_, f_n_*)⟩, where ⟨*r*_1_*, r*_2_*, r*_3_*, …, r_n_*⟩ is the sequence representation and ⟨*f*_1_*, f*_2_*, f*_3_*, …, f_n_*⟩ is the structure representation from Foldseek [59]. SaProt outputs the structure-aware vector representations (embeddings). Step 5: In constructing our training and test sets, we partitioned our dataset with respect to different train-versus-test sequence similarity thresholds via CD-HIT [22]. Step 6: We built a twohidden-layer perceptron that takes in the SaProt embedding of an RBP as input and outputs the host genus from among the ESKAPEE genera. Step 7: We evaluated our model’s performance. Graphics in this figure were taken from [1, 5, 3, 2, 4].

### 2.1 Data collection

We collected 20,941 phage genome sequences, alongside their respective protein sequences (if provided), that were uploaded to GenBank before October 2023. We used INPHARED [17], a pipeline for downloading phage sequences and host genus information from GenBank [11]. For phages where INPHARED did not return a host, we retrieved the isolation host information from their respective GenBank entries, if provided. Afterwards, we discarded phages with non-bacterial hosts, leaving a total of 17,790 phages across 273 host genera.

### 2.2 Receptor-binding protein identification

Among the 17,790 collected phage sequences, 16,627 already had genome annotations from GenBank. We ran the remaining 1,163 sequences without annotations through Prokka [48], which serves as a wrapper for a two-stage pipeline of calling Prodigal [30] for gene prediction and PHROG [57], a collection of viral protein families, for functional annotation.

With all the collected sequences annotated, we identified the receptor-binding proteins (RBPs). To this end, we matched the gene product annotations against a regular expression that we proposed in our previous work [23] based on the pattern given by Boeckaerts et al. [13]. Since this pattern also captures RBP-related annotations that are not RBPs per se (e.g., tailspike adaptor proteins), we referred to the exclusion list provided by Boeckaerts et al. [13] to filter out these cases.

Moreover, in view of the possibility that some proteins tagged as hypothetical might be RBPs, we converted these proteins into embeddings using the protein language model ProtBert [21] and then passed the resulting embeddings as input to the extreme gradient boosting model PhageRBPdetect [13] to predict whether the proteins are RBPs.

We subsequently discarded RBPs with outlying lengths, i.e., those with lengths outside the interval [*Q*_1_ − 1.5 · *IQR, Q*_3_ + 1.5 · *IQR*], where *Q*_1_, *IQR*, and *Q*_3_ refer to the first quartile, interquartile range, and third quartile of the RBP lengths, respectively. Finally, we removed duplicate RBPs using CD-HIT [22], a tool for clustering protein sequences.

### 2.3 Protein structure prediction

We fed the RBP sequences to ColabFold [43], a protein structure prediction tool that accelerates the inference time of AlphaFold 2 [33] by building multiple sequence alignments using MMseqs2 [51] instead of JackHMMer [32]. ColabFold predicts the *x*-, *y*-, and *z*-coordinates of all the heavy atoms (carbon, hydrogen, nitrogen, oxygen, and sulfur) of a given protein. Table A1 lists the parameters at which we ran ColabFold.

Our dataset consists of the predicted structures of 7,627 RBPs from 3,350 phages targeting hosts from the ESKAPEE genera (Table 1). The distributions of the RBP lengths and ColabFold’s confidence scores per genus are plotted in Figures 2 and 3. We computed the confidence score for each protein by averaging the predicted local distance difference test (pLDDT) scores across its residues; the pLDDT score is a superposition-free estimate of the agreement between a predicted structure and a reference structure [33]. For each of the ESKAPEE genera, the mean confidence score is above 77% and the median is above 80%.

**Figure 2:**
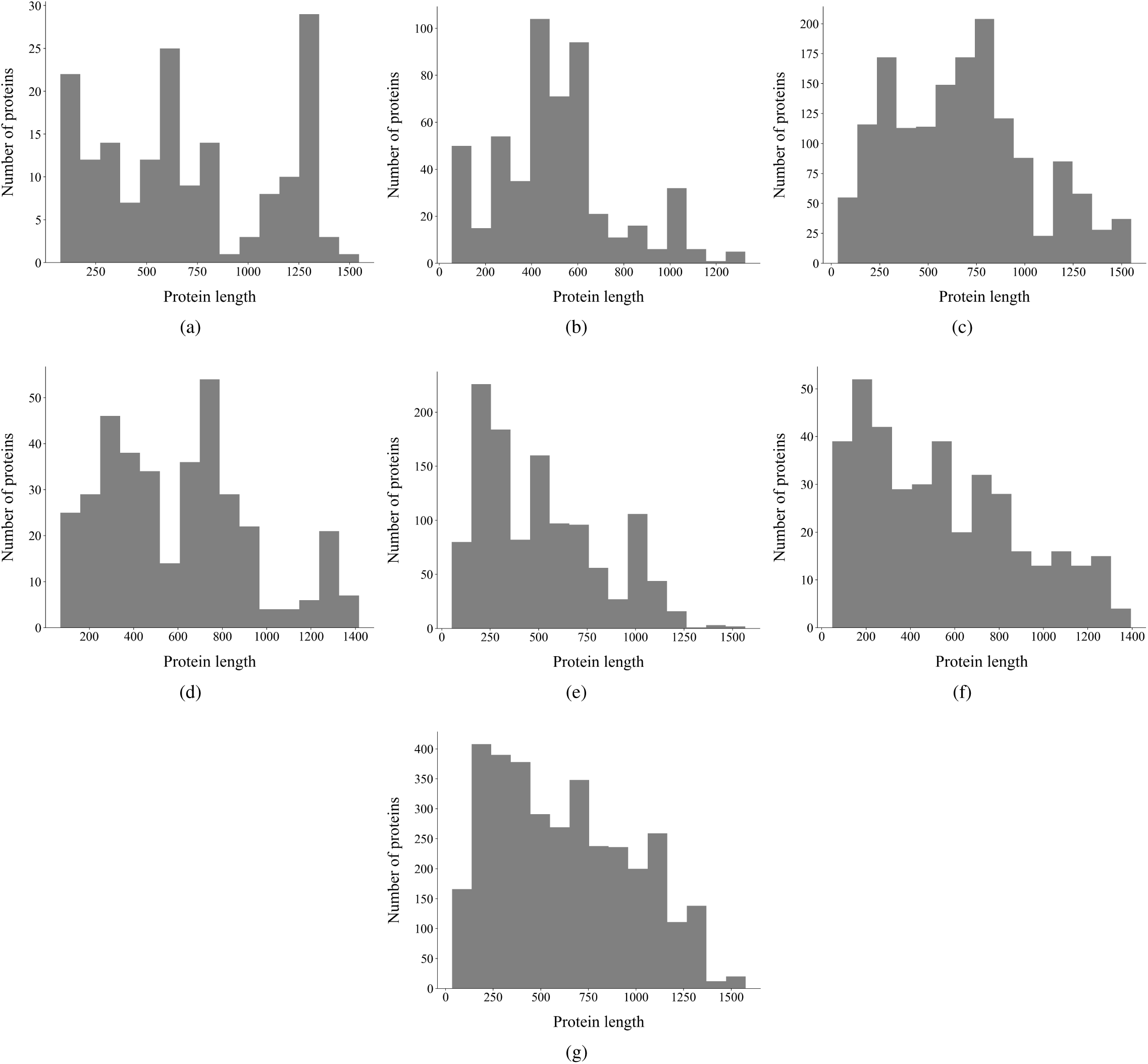
Per-genus distribution of the receptor-binding protein lengths (number of amino acids): (a) *Enterococcus*, (b) *Staphylococcus*, (c) *Klebsiella*, (d) *Acinetobacter*, (e) *Pseudomonas*, (f) *Enterobacter*, and (g) *Escherichia*.

**Figure 3:**
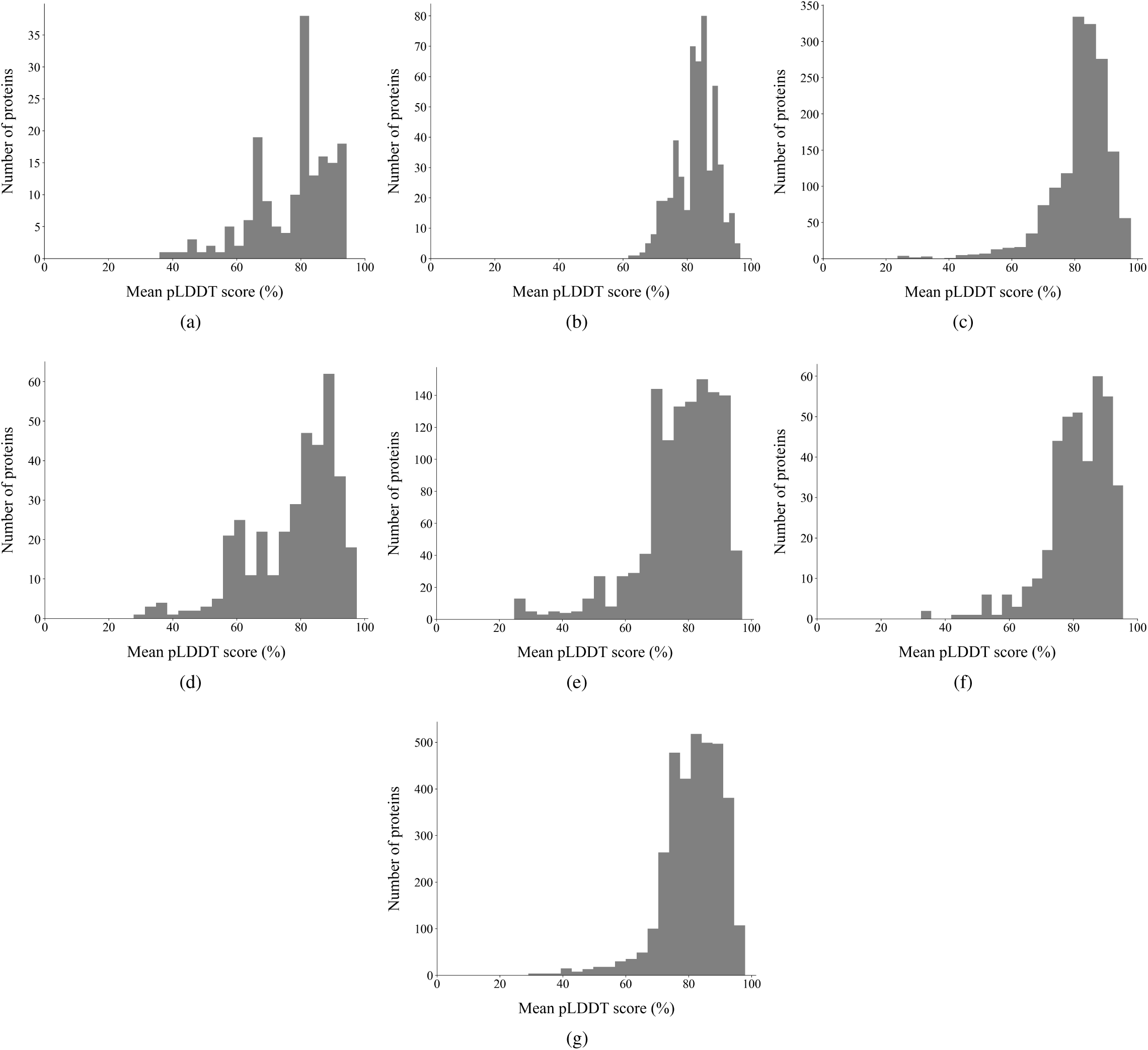
Per-genus distribution of ColabFold’s confidence scores in its predicted protein structures: (a) *Enterococcus*, (b) *Staphylococcus*, (c) *Klebsiella*, (d) *Acinetobacter*, (e) *Pseudomonas*, (f) *Enterobacter*, and (g) *Escherichia*. The confidence score for each protein was calculated by taking the mean predicted local distance difference test (pLDDT) scores across its residues.

**Table 1:** Per-genus dataset characteristics.

| Genus | Count | RBP length (a.a.) |  | Mean pLDDT score |  |
| --- | --- | --- | --- | --- | --- |
|  |  | Mean | Median | Mean | Median |
| *Enterococcus* | 170 | 704 | 620 | 77.57% | 80.87% |
| *Staphylococcus* | 521 | 510 | 481 | 82.74% | 83.24% |
| *Klebsiella* | 1,535 | 671 | 658 | 82.00% | 83.48% |
| *Acinetobacter* | 369 | 599 | 607 | 77.80% | 81.53% |
| *Pseudomonas* | 1,180 | 509 | 477 | 77.48% | 79.50% |
| *Enterobacter* | 388 | 545 | 512 | 81.26% | 82.24% |
| *Escherichia* | 3,464 | 622 | 581 | 81.39% | 82.42% |

We also ran our pipeline on phages that target other host genera, thereby expanding our dataset of RBPs and their predicted structures to 19,081 RBPs from 8,525 phages across 238 host genera.

Figure A1a shows the distribution of the RBP lengths; the mean length is 565 amino acids, and the median is 507 amino acids. Figure A1b shows the distribution of ColabFold’s confidence scores in its predicted protein structures. The mean confidence score across all the proteins in our dataset is 78.45%, and the median is 81.69%.

### 2.4 Structure-aware embedding generation

In order to generate the vector representations (embeddings) of the RBPs, we employed SaProt, a structure-aware protein language model that adopts the architecture of ESM-2 [40] but with the embedding layer modified in view of its structure-aware alphabet. SaProt’s alphabet attempts to capture both sequence and structure information by taking the Cartesian product of the set of all possible residues and Foldseek’s [59] alphabet. Foldseek’s alphabet consists of 20 letters that describe the tertiary interaction between residues and their nearest neighbors; this alphabet was learned via a vector quantized variational autoencoder [58]. Formally, suppose we are given a protein of length *n*. Its sequence representation is ⟨*r*_1_*, r*_2_*, r*_3_*, …, r_n_*⟩, where *r_i_* is its *i*^th^ residue. Foldseek encodes its structure information into an *n*-letter string ⟨*f*_1_*, f*_2_*, f*_3_*, …, f_n_*⟩. The input to SaProt is ⟨(*r*_1_*, f*_1_), (*r*_2_*, f*_2_), (*r*_3_*, f*_3_) *, …,* (*r_n_, f_n_*)⟩. In order to obtain a fixed-length embedding, we averaged the last layer’s hidden states over the sequence length, resulting in a 1,280-long dense vector representation (structure-aware embedding) for each RBP. We used the 650-million-parameter version of SaProt that was pretrained on 40 million structures from AlphaFold DB [61].

### 2.5 Sequence similarity-based training and test set construction

We investigated model performance at different train-versus-test sequence similarity thresholds. A lower train-versus-test sequence similarity indicates that the sequences in the training set are more dissimilar to those in the test set.

To this end, we used CD-HIT [22] to cluster the RBPs at a similarity threshold *s* and assign a representative sequence per cluster. Let *R* be the set of representative sequences with class labels among the ESKAPEE genera. We split *R* into two sets *D*_1_ and *D*_2_, with 70% of the sequences assigned to *D*_1_ and the remaining 30% to *D*_2_; this splitting was stratified based on the class sizes. Clusters with representative sequences in *D*_1_ were assigned to the training set, while those with representative sequences in *D*_2_ were assigned to the test set. We also randomly sampled RBPs with class labels outside the ESKAPEE genera and added these to our test set. The rationale is for the performance evaluation to reflect scenarios where our model is fed with an RBP from a phage infecting a host outside our genera of interest.

In order to mitigate class imbalance (Table A2), we performed data augmentation via SMOTETomek [10]. Our training and test set statistics are reported in Table 2; the per-class breakdown is given in Table A2.

**Table 2:** Training and test set statistics.

| Sequence similarity $s$ | Training set size | | Test set size |
| --- | --- | --- | --- |
|  | Before SMOTE-Tomek | After SMOTE-Tomek |  |
| 100% | 5,338 | 16,942 | 2,340 |
| 80% | 4,865 | 14,822 | 2,822 |
| 60% | 4,523 | 13,099 | 3,153 |
| 40% | 4,465 | 11,662 | 3,203 |

### 2.6 Classifier building

We defined phage-host interaction prediction as a multiclass classification task where the input is the SaProt embedding of a given RBP and the output is one of ESKAPEE host genera.

To this end, we built a multilayer perceptron with two hidden layers (Figure 4). The first and second hidden layers have 160 (one-eighth of the size of the input layer) and 80 neurons, respectively. We implemented *L*_2_ regularization and applied dropout with a rate of 0.2 after the first hidden layer. We employed rectified linear unit as the activation function, softmax as the output function, and crossentropy as the loss function. We trained the model for 200 epochs with Adam [34] as the optimizer; other training hyperparameters are listed in Table 3. Hyperparameter tuning was performed using a validation set constructed by setting aside 10% of our training set. We call the resulting model PHIStruct.

**Figure 4:**
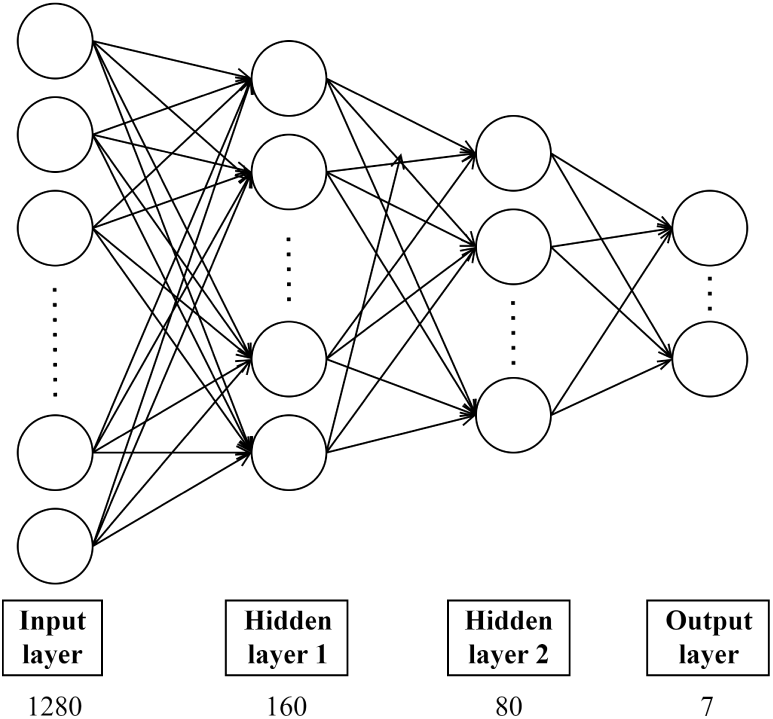
PHIStruct classifier architecture. The number below the label of each layer denotes the size of that layer.

**Table 3:**
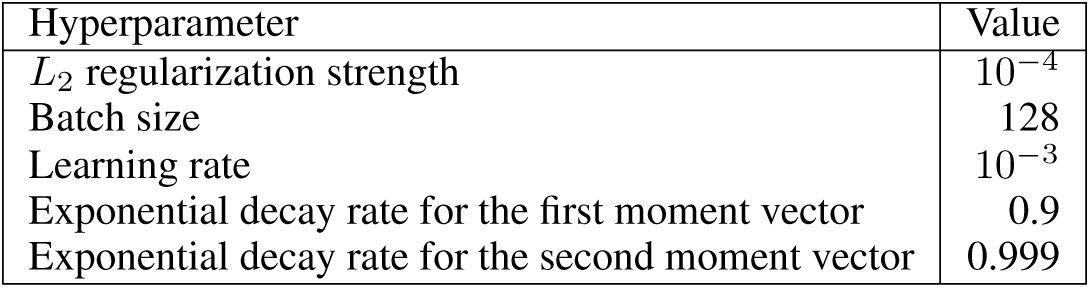
PHIStruct training hyperparameters.

### 2.7 Performance evaluation

We evaluated the performance of our model in terms of class-averaged metrics: macro-precision, macro-recall, and macro-F1. To account for our model’s confidence in its prediction, we parameterized these metrics based on a confidence threshold *k*. Let *p*_1_ and *p*_2_ be the class probabilities for the model’s highest-ranked and its second-highest-ranked predictions, respectively. The input is classified under the highest-ranked prediction only if *p*_1_ − *p*_2_ ≥ *k*. In this regard, higher values of *k* prioritize precision, whereas lower values of *k* prioritize recall. We also parameterized the metrics based on the maximum train-versus-test sequence similarity *s*.

Let *C* be the set of ESKAPEE class labels, and let *TP_c,k,s_*, *TN_c,k,s_*, *FP_c,k,s_*, and *FN_c,k,s_* denote the number of true positive, true negative, false positive, and false negative classifications for class *c* ∈ *C*, respectively, at confidence threshold *k* and maximum train-versus-test sequence similarity *s*. We define our metrics formally in Equations 1 to 3.

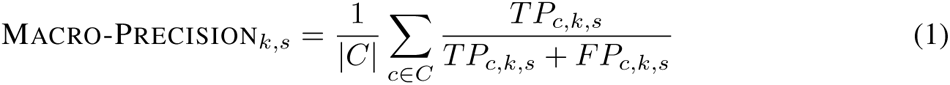

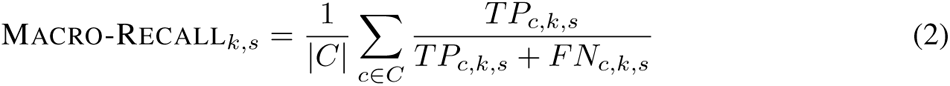

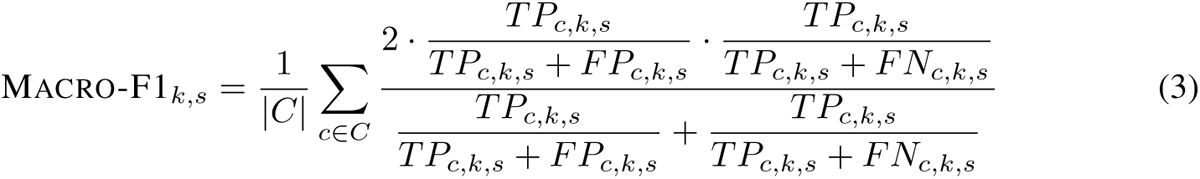

## 3 Results and discussion

### 3.1 At lower train-versus-test sequence similarity, PHIStruct outperforms state-of-the-art machine learning tools that take in receptor-binding proteins as input, as well as BLASTp

We benchmarked our tool, PHIStruct, against state-of-the-art machine learning tools that also map receptor-binding proteins to host bacteria: Boeckaerts et al.’s tool [12], PHIEmbed [23], and Badam and Rao’s tool [9]. Boeckaerts et al.’s tool [12] is a random forest model that takes in manually feature-engineered genomic and protein sequence properties. Our previous work, PHIEmbed, is a random forest model that takes in ProtT5 [21] embeddings. Badam and Rao’s tool [9] is a multilayer perceptron that takes in ESM-1b [47] embeddings. To ensure a fair comparison, we retrained them on our dataset. Additionally, we benchmarked against BLASTp, which we ran at an E-value of 0.05 and with BLOSUM62 as the scoring matrix; the class label of the reported top hit was taken as the predicted host genus. We evaluated the tools’ performance across maximum train-versus-test sequence similarity thresholds *s* = 40% to 100% in steps of 20% and across confidence thresholds *k* = 0% to 90% in steps of 10%.

Our experiments showed that PHIStruct presents improvements over these tools, especially at low sequence similarity settings. Its performance gains were most pronounced at *s* = 40% (Figure 5), where it registered a maximum recall of 63.09%, thus achieving a margin of 11% over PHIEmbed, 5% over BLASTp, and 4% over Badam and Rao’s tool [9] and performing competitively with Boeckaerts et al.’s tool [12]. It obtained a maximum precision of 69.43%, outperforming Badam and Rao’s tool [9] by 11% and BLASTp by 16%. Moreover, it recorded the highest maximum F1 at 57.67%, outperforming Boeckaerts et al.’s tool [12] by 2%, BLASTp by 6%, Badam and Rao’s tool [9] by 8%, and PHIEmbed by 14%.

**Figure 5:**
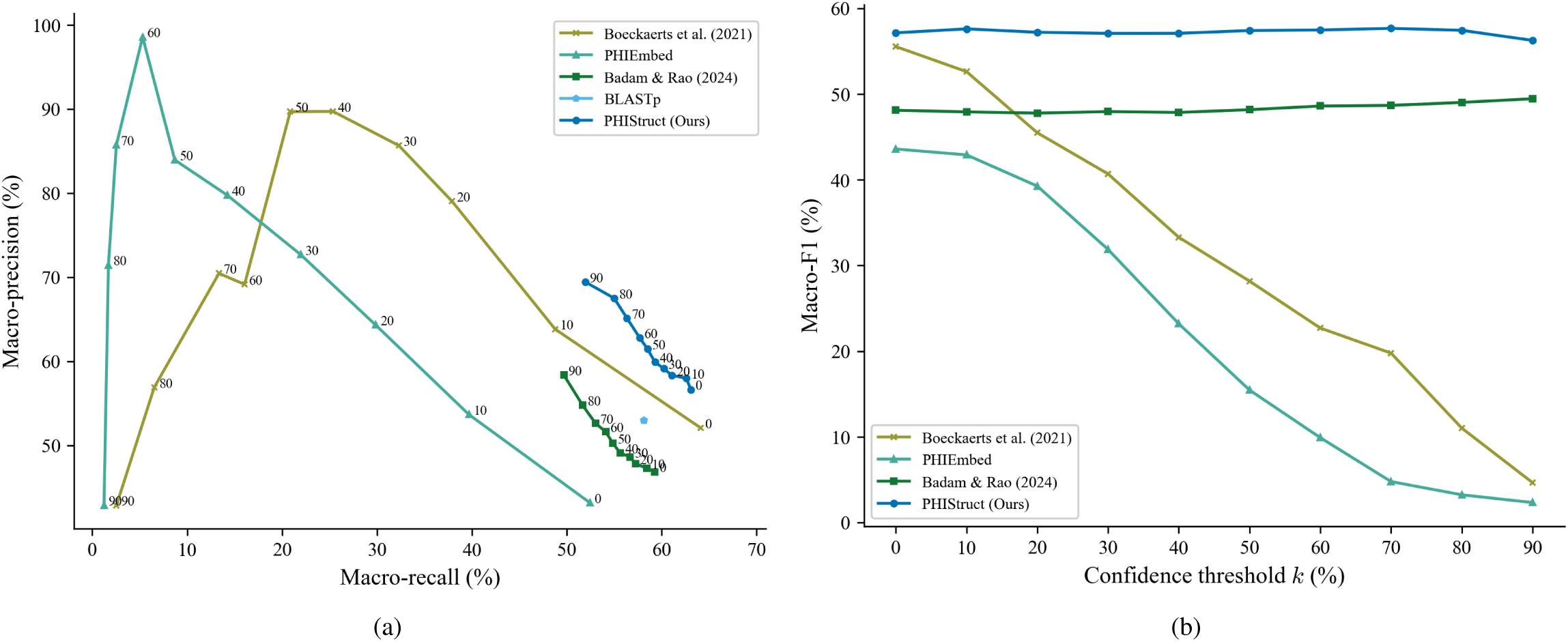
Comparison of the performance of PHIStruct with BLASTp and state-of-the-art machine learning tools that map receptor-binding proteins to host bacteria. The maximum train-versus-test sequence similarity is set to *s* = 40%. (a) Precision-recall curves. The label of each point denotes the confidence threshold *k* (%) at which the performance was measured. (b) F1 scores. Higher values of *k* prioritize precision over recall, whereas lower values prioritize recall.

**Figure 6:**
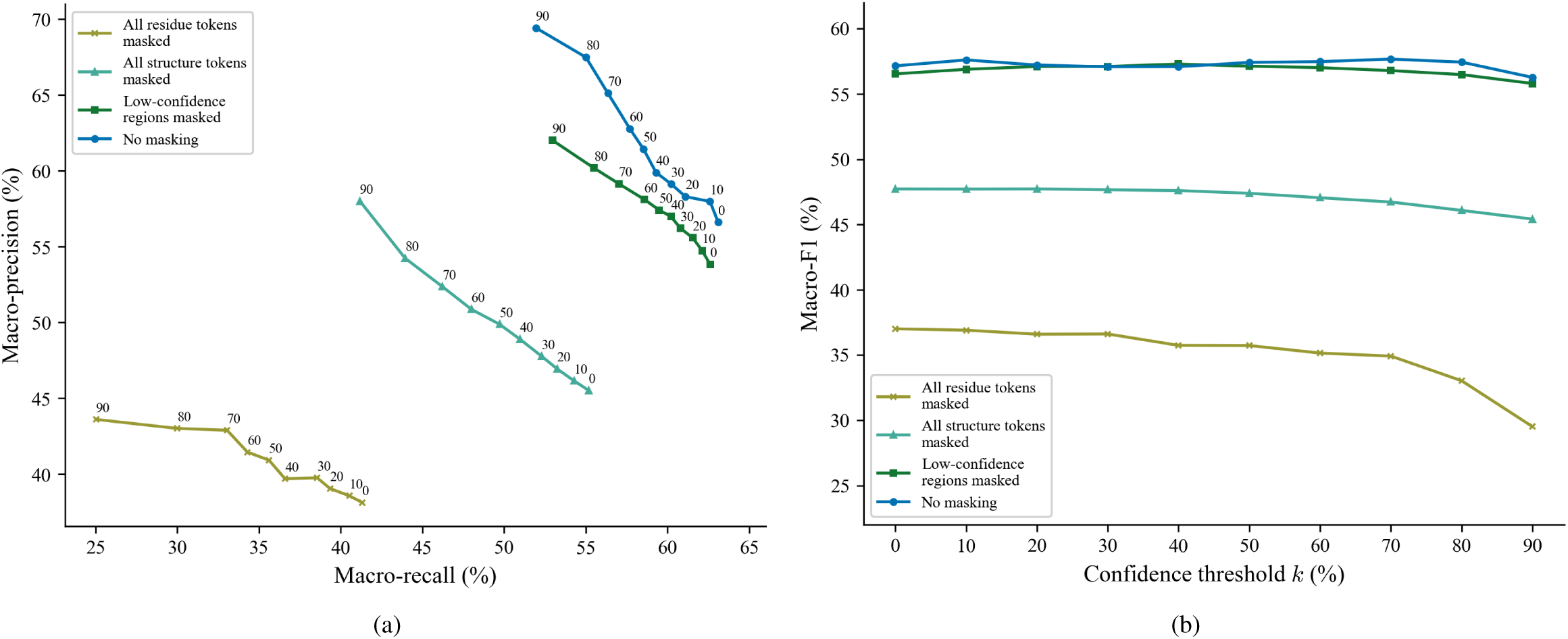
Comparison of the performance of different masking strategies for inputting proteins to SaProt. The maximum train-versus-test sequence similarity is set to *s* = 40%. (a) Precision-recall curves. The label of each point denotes the confidence threshold *k* (%) at which the performance was measured. (b) F1 scores. Higher values of *k* prioritize precision over recall, whereas lower values prioritize recall.

Although Boeckaerts et al.’s tool [12] and PHIEmbed had higher maximum precision scores, these were accompanied by a marked loss in recall. This trade-off was less pronounced with PHIStruct, as reflected in the precision-recall curves (Figures 5a and Figures A2a to A4a) and F1 scores (Figure 5b and Figures A2b to A4b). Across all the tested sequence similarity thresholds, the decrease in its F1 from *k* = 0% to 90% was under 2%. Furthermore, at *k >* 50%, it outperformed the machine learning tool with the next-highest F1 by 7% to 9% and BLASTp by 5% to 6% in terms of F1. These results highlight our tool’s ability to make high-confidence predictions without heavily compromising between precision and recall.

PHIStruct also performed competitively at higher sequence similarity thresholds. At *s* = 60%, it recorded a maximum recall of 65.83%, maximum precision of 71.77%, and maximum F1 of 62.95%, outperforming the other machine learning tools in terms of maximum recall and F1 (Figure A2). At *s* = 80%, it obtained a maximum recall of 67.08%, maximum precision of 69.10%, and maximum F1 of 63.73%, placing its maximum recall within 1.6% of the tool with the highest maximum recall (Figure A3). At *s* = 100%, it reported a maximum recall of 81.69%, maximum precision of 87.40%, and maximum F1 of 81.12%, placing its maximum recall within 0.4% of the top-performing machine learning tool and its maximum F1 within 0.1% (Figure A4).

These results demonstrate the utility of PHIStruct for improved prediction of phage-host pairs, especially in use cases where phages of interest have receptor-binding proteins with low sequence similarity to those of known phages. The per-genus results are reported in Tables A3 to A5.

### 3.2 Sequence information and structure information are complementary in improving performance, especially at lower train-versus-test sequence similarity

As discussed in Section “Structure-aware embedding generation”, prior to the generation of the structure-aware embedding, SaProt requires a protein to be input as the sequence ⟨(*r*_1_*, f*_1_), (*r*_2_*, f*_2_), (*r*_3_*, f*_3_) *, …,* (*r_n_, f_n_*)⟩, where *r_i_* is the *i*^th^ residue and *f_i_* is the corresponding structure (Foldseek) token.

To investigate the impact of incorporating structure information in this input step, we benchmarked different masking strategies: *(i)* masking all residue tokens, i.e., replacing all (*r_i_, f_i_*) with (#*, f_i_*), where # is a special token denoting the mask, *(ii)* masking all structure tokens, i.e., replacing all (*r_i_, f_i_*) with (*r_i_,* #), *(iii)* masking the structure tokens of low-confidence regions, and *(iv)* not masking any token (which we employed in PHIStruct). For the third masking strategy, low-confidence regions are defined as those having a pLDDT score below 70%, following the threshold in previous studies [54, 60].

Our experiments showed that including both sequence and structure tokens improves performance, especially at lower train-versus-test sequence similarity thresholds. At *s* = 40%, the highest performance across all three metrics was obtained by not masking any token. Masking all the structure tokens resulted in an 8%, 11%, and 10% decrease in maximum recall, precision, and F1, respectively, whereas masking all the residue tokens resulted in a 22%, 26%, and 21% decrease in these aforementioned metrics. Not masking any token also returned the highest performance at *s* = 60% and 80% (Figures A5 and A6). At *s* = 100%, the highest performance was achieved by masking the structure tokens of low-confidence regions (Figure A7).

These results suggest that, as the sequence similarity between an RBP of interest and known RBPs decreases, the model benefits from complementing sequence information with structure-informed signals.

### 3.3 Differences in the receptor-binding protein embeddings in relation to the host genus may be contributing to PHIStruct’s discriminative power

To investigate the differences in the embeddings in relation to the host genus, we paired the RBPs such that the sequence similarity between the RBPs in each pair is below 40%. We then computed the cosine distance between the SaProt embeddings of the RBPs in each pair. Afterwards, for every genus, we constructed two groups, each with 500 randomly sampled RBP pairs. In the first group, both RBPs in each pair target the same genus of interest. In the second group, one of the RBPs in each pair targets the genus of interest, while the other RBP targets a different genus. Performing a Mann-Whitney U test showed that, for all but two genera, the distribution of the cosine distance values between these two groups is statistically significant at a *p*-value cutoff of 0.05 (Table A7).

It is possible that PHIStruct may be capturing these differences in the structure-aware embeddings in relation to the host genus, thereby contributing to the model’s discriminative power.

### 3.4 PHIStruct’s use of structure-aware embeddings outperforms using sequence-only embeddings, especially at lower train-versus-test sequence similarity

To investigate the impact of the protein representation (i.e., structure-aware versus sequence-only embeddings), we benchmarked PHIStruct against multilayer perceptron models that take in embeddings from ProtT5 [21], ESM-1b [47], and ESM-2 [40], state-of-the-art sequence-only protein language models used in existing phage-host interaction prediction tools [23, 9, 14]. The specific model versions used in our benchmarking experiments are given in Table A8. The downstream multilayer perceptron models share the same architecture as that of PHIStruct: their first and second hidden layers have *<u>^L^</u>* and *<u>^L^</u>* neurons, respectively, where *L* is the length of the protein embedding.

Our experiments showed that our use of structure-aware embeddings presents improvements over using sequence-only embeddings, especially at lower train-versus-test sequence similarity thresholds. At *s* = 40%, PHIStruct’s margins over the model with the next-highest performance (i.e., the model using ProtT5) were 7% in terms of maximum precision and 2% in terms of maximum recall and F1 (Figure 7). At *s* = 60%, it outperformed the model with the next-highest maximum recall (i.e., the model using ProtT5) by 2%, the model with the next-highest maximum precision (i.e., the model using ESM-1b) by 3%, and the model with the next-highest maximum F1 (i.e., the model using ProtT5) by 4% (Figure A8). At *s* = 80%, it obtained the highest maximum recall (Figure A9). At *s* = 100%, it registered the highest maximum precision and F1 (Figure A10).

**Figure 7:**
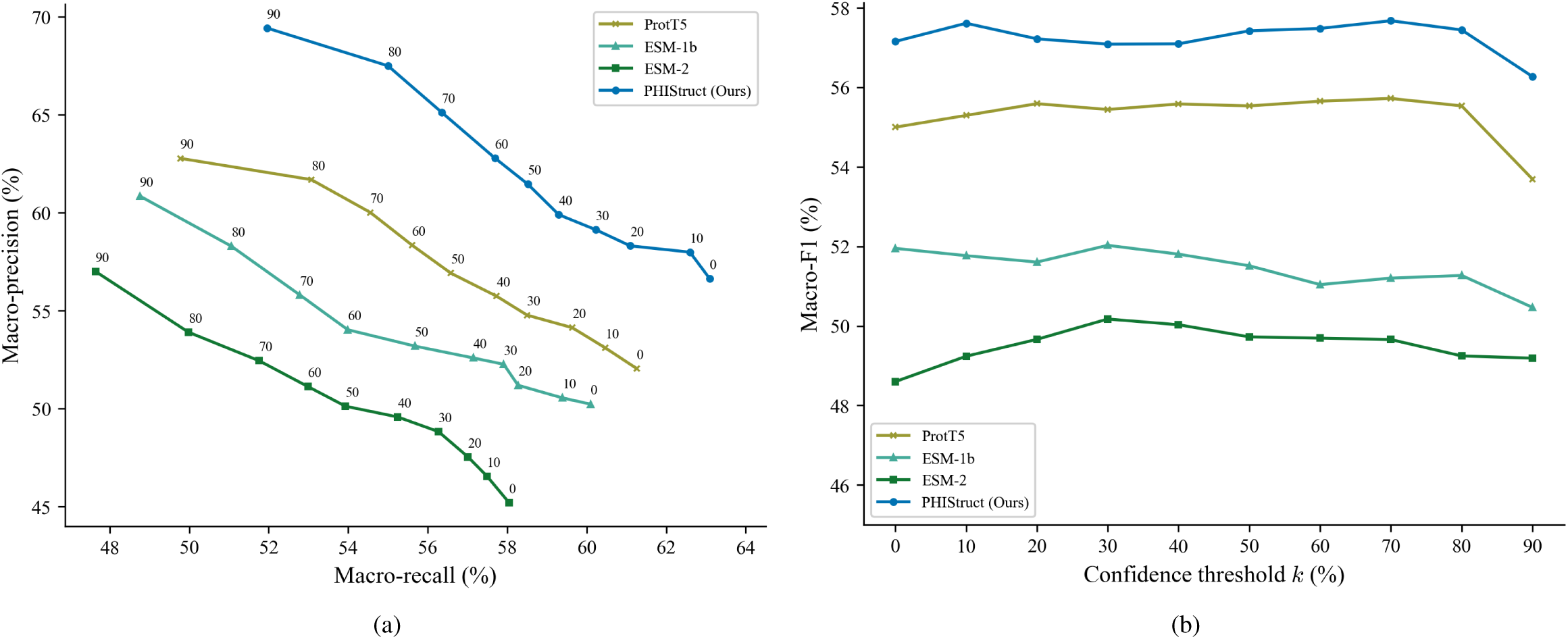
Comparison of the performance of PHIStruct with same-architecture multilayer perceptron models that take in sequence-only embeddings. The maximum train-versus-test sequence similarity is set to *s* = 40%. (a) Precision-recall curves. The label of each point denotes the confidence threshold *k* (%) at which the performance was measured. (b) F1 scores. Higher values of *k* prioritize precision over recall, whereas lower values prioritize recall.

### 3.5 PHIStruct’s use of SaProt embeddings outperforms using other structure-aware protein embeddings, especially at lower train-versus-test sequence similarity

To further investigate the impact of the protein representation (i.e., SaProt embeddings versus other structure-aware embeddings), we benchmarked PHIStruct against multilayer perceptron models that share the same architecture as that of PHIStruct but take in embeddings from other structure-aware protein language models: ProstT5 [28] and PST [16]. ProstT5 adopts the architecture of ProtT5 [21] and is fine-tuned to convert bidirectionally between protein sequence and structure. PST adopts the architecture of ESM-2 [40] and augments each self-attention block with a two-layer graph isomorphism network [65] that serves as a protein structure extractor module. The specific model versions used in our benchmarking experiments are given in Table A8.

Our experiments showed that our use of SaProt embeddings presents improvements over using other structure-aware embeddings, especially at lower train-versus-test sequence similarity thresholds. From *s* = 40% to 80%, PHIStruct achieved the highest performance across all three metrics (Figures 8, A11, and A12). In particular, its performance gains were most pronounced at *s* = 40%, where it outperformed the model with the next-highest performance (i.e., the model using PST) by 5% in terms of maximum recall, 13% in terms of maximum precision, and 8% in terms of maximum F1 (Figure 8). At *s* = 100%, it obtained the highest maximum precision and F1, and its maximum recall was within 1.2% of the model with the highest maximum recall (Figure A13).

**Figure 8:**
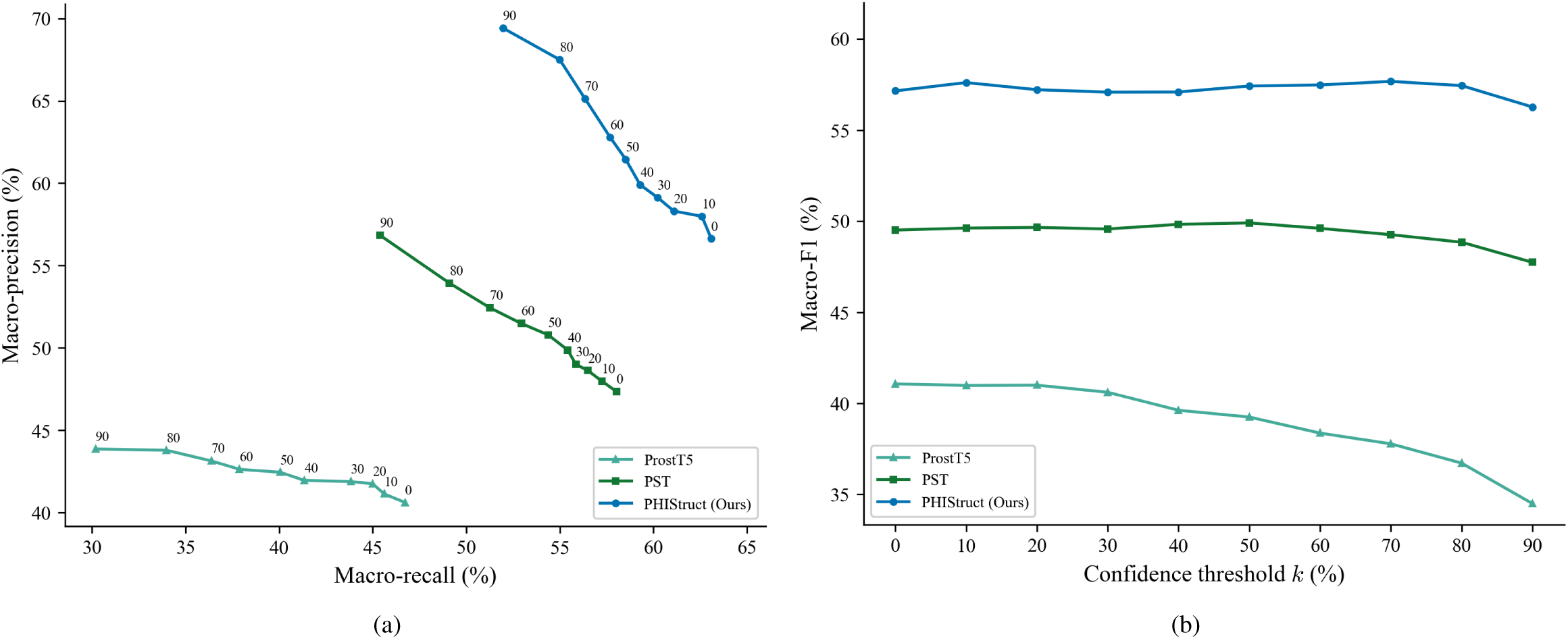
Comparison of the performance of PHIStruct with same-architecture multilayer perceptron models that take in structure-aware protein embeddings other than SaProt. The maximum trainversus-test sequence similarity is set to *s* = 40%. (a) Precision-recall curves. The label of each point denotes the confidence threshold *k* (%) at which the performance was measured. (b) F1 scores. Higher values of *k* prioritize precision over recall, whereas lower values prioritize recall.

### 3.6 Room for improvement

We performed error analysis on PHIStruct’s predictions and found that, at lower train-versus-test sequence similarity thresholds, it tends to misclassify phages that infect *Enterobacter* (and *Klebsiella*, albeit to a lesser degree) as infecting *Escherichia* (Figure 9). This could possibly be related to these three genera belong to the same family, Enterobacteriaceae. The four remaining ESKAPEE genera each belong to different families.

**Figure 9:**
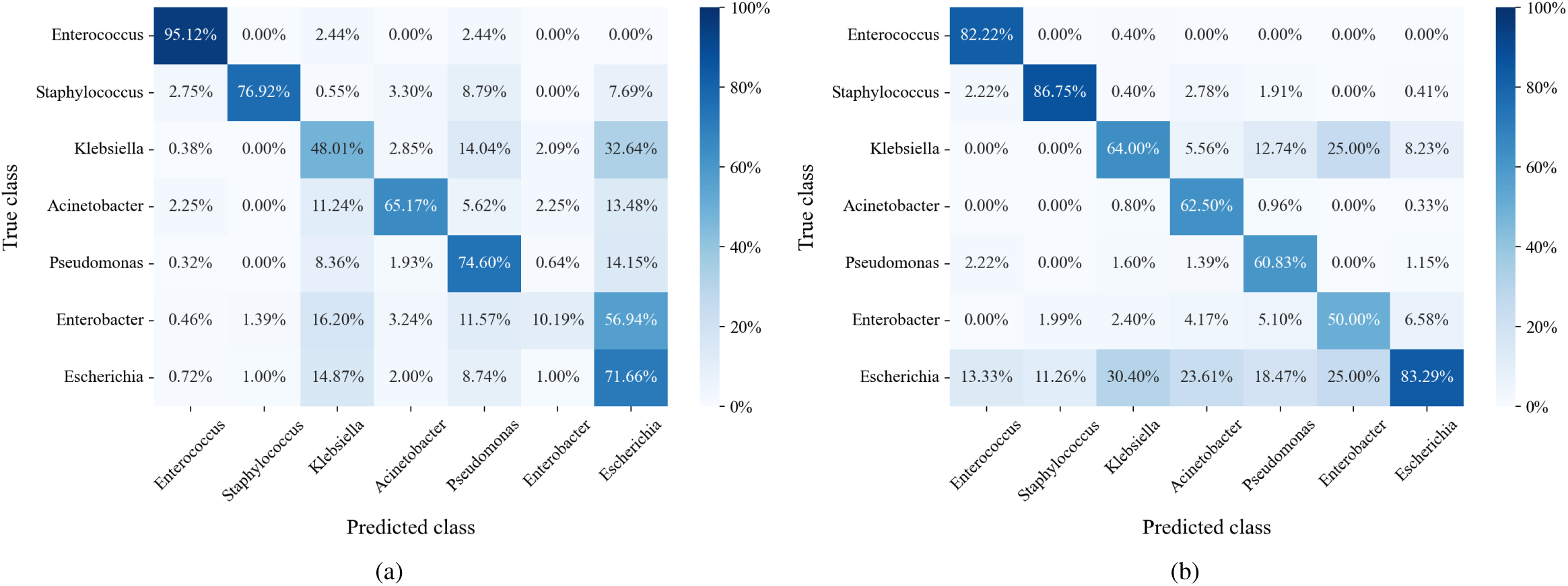
Confusion matrices at maximum train-versus-test sequence similarity *s* = 40%. (a) Confusion matrix at confidence threshold *k* = 0%, normalized over the true class labels. The main diagonal reflects the per-class recall. (b) Confusion matrix at *k* = 90%, normalized over the predicted class labels. The main diagonal reflects the per-class precision. Lower values of *k* prioritize recall over precision, whereas higher values prioritize precision.

In this regard, possible directions for future work include investigating biological markers that can more strongly distinguish phage-host specificity at lower taxonomic levels. It may also be helpful to develop computational approaches that are explicitly trained to discriminate between biological signals at finer taxonomic resolutions.

## 4 Conclusion

In this paper, we presented PHIStruct, a deep learning model that capitalizes on structure-aware embeddings of receptor-binding proteins for predicting phage-host interaction. To this end, we predicted the protein structures via ColabFold and generated the structure-aware embeddings via SaProt. Our experiments showed that our proposed approach presents improvements over stateof-the-art methods, especially as the sequence similarity between the training and test set entries decreases. These results highlight the applicability of PHIStruct in improving the prediction of phage-host pairs, especially in settings where newly discovered phages have receptor-binding proteins with low sequence similarity to those of known phages.

With recent investigations showing an association between tailspike proteins and bacterial polysaccharide receptors [66], future directions include jointly incorporating these proteins, alongside other phage-encoded and host-encoded proteins involved in different stages of the phage infection process.

## 5 Data availability statement

The data and source code for our experiments and analyses are available at https://github.com/bioinfodlsu/PHIStruct. Our dataset of ColabFold-predicted structures of receptor-binding proteins is available at https://doi.org/10.5281/zenodo.11202338.

## Acknowledgments

The authors thank Dr. Ann Franchesca B. Laguna for her advice on the methodology. This research was partly funded by the Department of Science and Technology Philippine Council for Health Research and Development (DOST-PCHRD) under the e-Asia JRP 2021 Alternative therapeutics to tackle AMR pathogens (ATTACK-AMR) program. This research was supported with Cloud TPUs from Google’s TPU Research Cloud (TRC) and with computing resources from the Machine Learning eResearch Platform (MLeRP) of Monash University, University of Queensland, and Queensland Cyber Infrastructure Foundation Ltd.

## A Appendix

**Figure A1:**
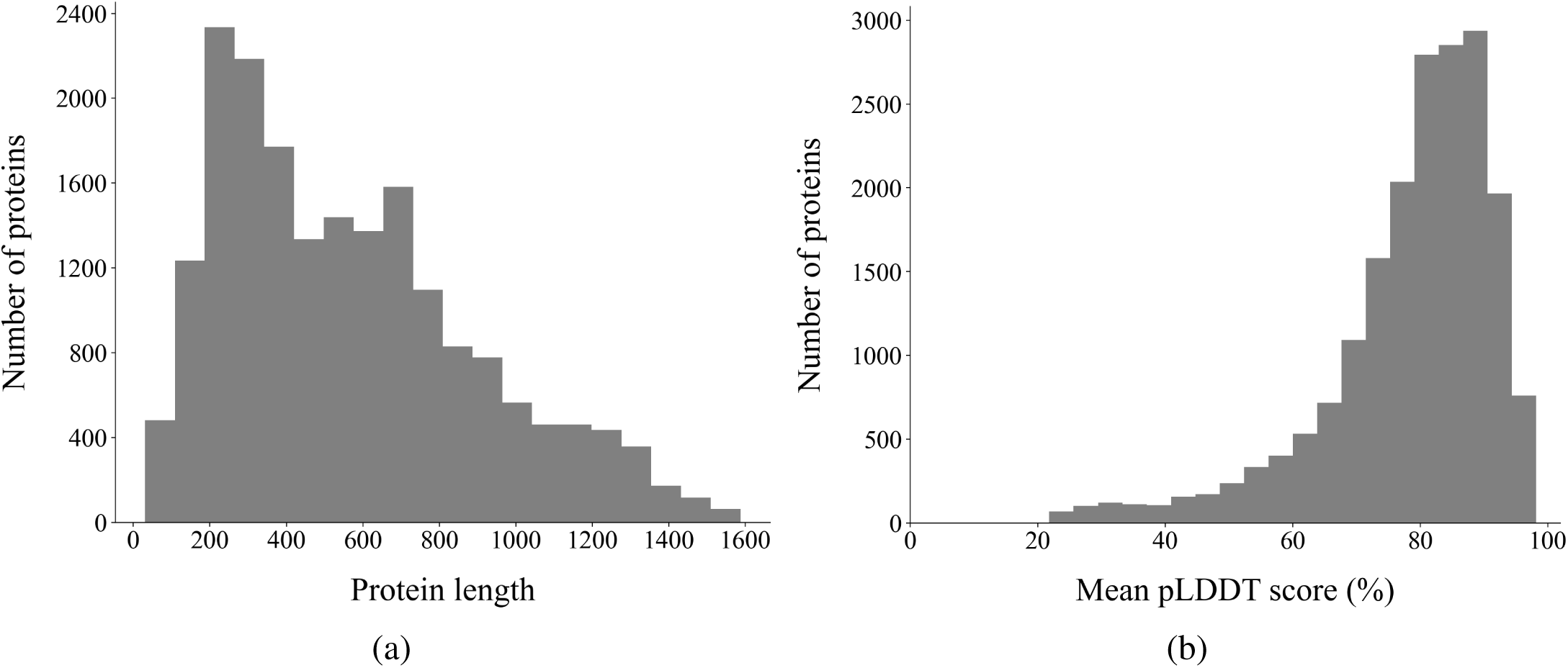
Dataset characteristics. (a) Distribution of receptor-binding protein lengths (number of amino acids). (b) Distribution of ColabFold’s confidence scores in its predicted protein structures. The confidence score for each protein was calculated by taking the mean predicted local distance difference test (pLDDT) scores across its residues.

**Figure A2:**
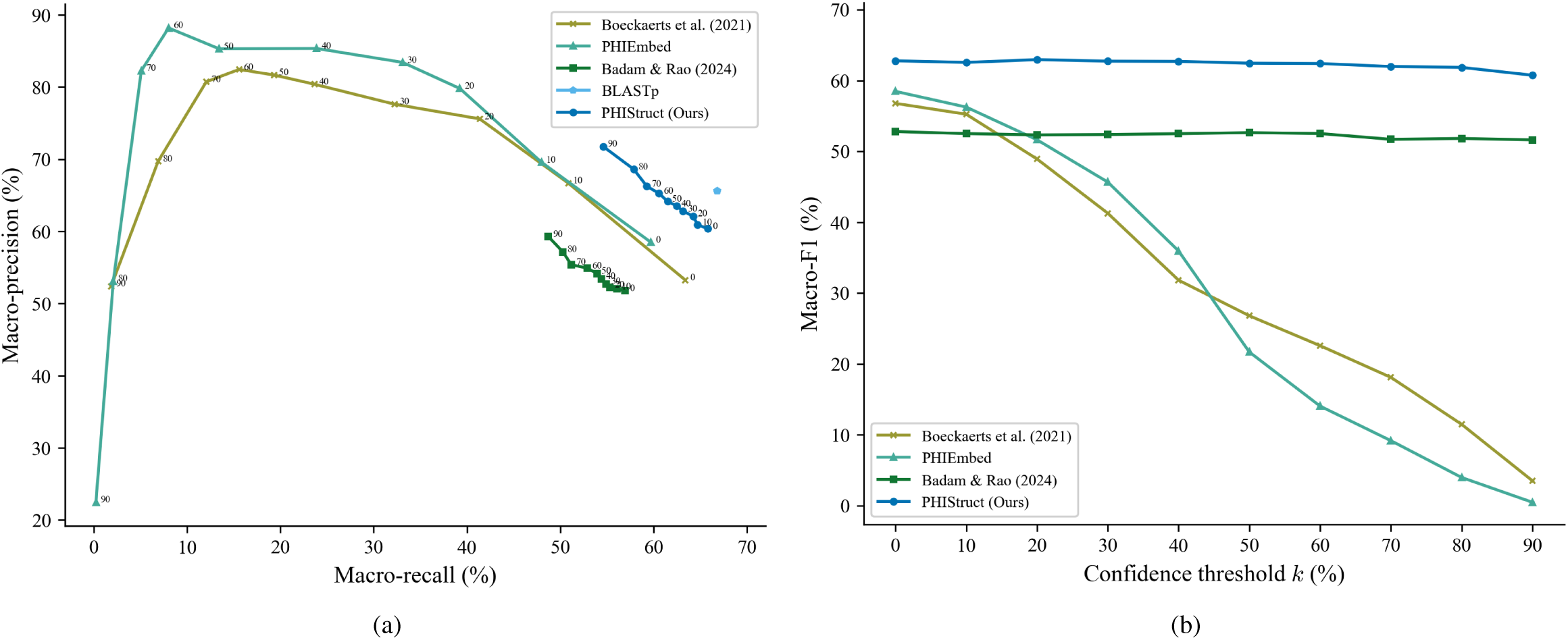
Comparison of the performance of PHIStruct with BLASTp and state-of-the-art machine learning tools that map receptor-binding proteins to host bacteria. The maximum train-versus-test sequence similarity is set to *s* = 60%. (a) Precision-recall curves. The label of each point denotes the confidence threshold *k* (%) at which the performance was measured. (b) F1 scores. Higher values of *k* prioritize precision over recall, whereas lower values prioritize recall.

**Figure A3:**
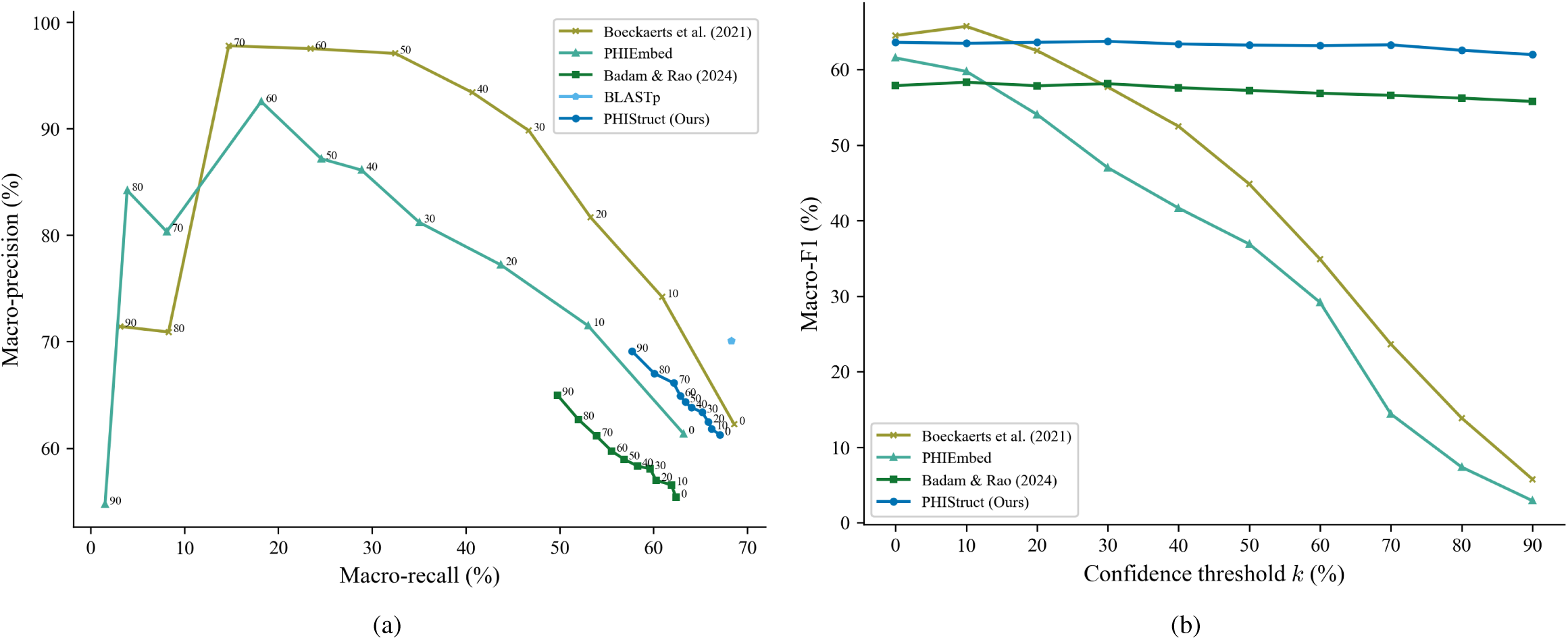
Comparison of the performance of PHIStruct with BLASTp and state-of-the-art machine learning tools that map receptor-binding proteins to host bacteria. The maximum train-versus-test sequence similarity is set to *s* = 80%. (a) Precision-recall curves. The label of each point denotes the confidence threshold *k* (%) at which the performance was measured. (b) F1 scores. Higher values of *k* prioritize precision over recall, whereas lower values prioritize recall.

**Figure A4:**
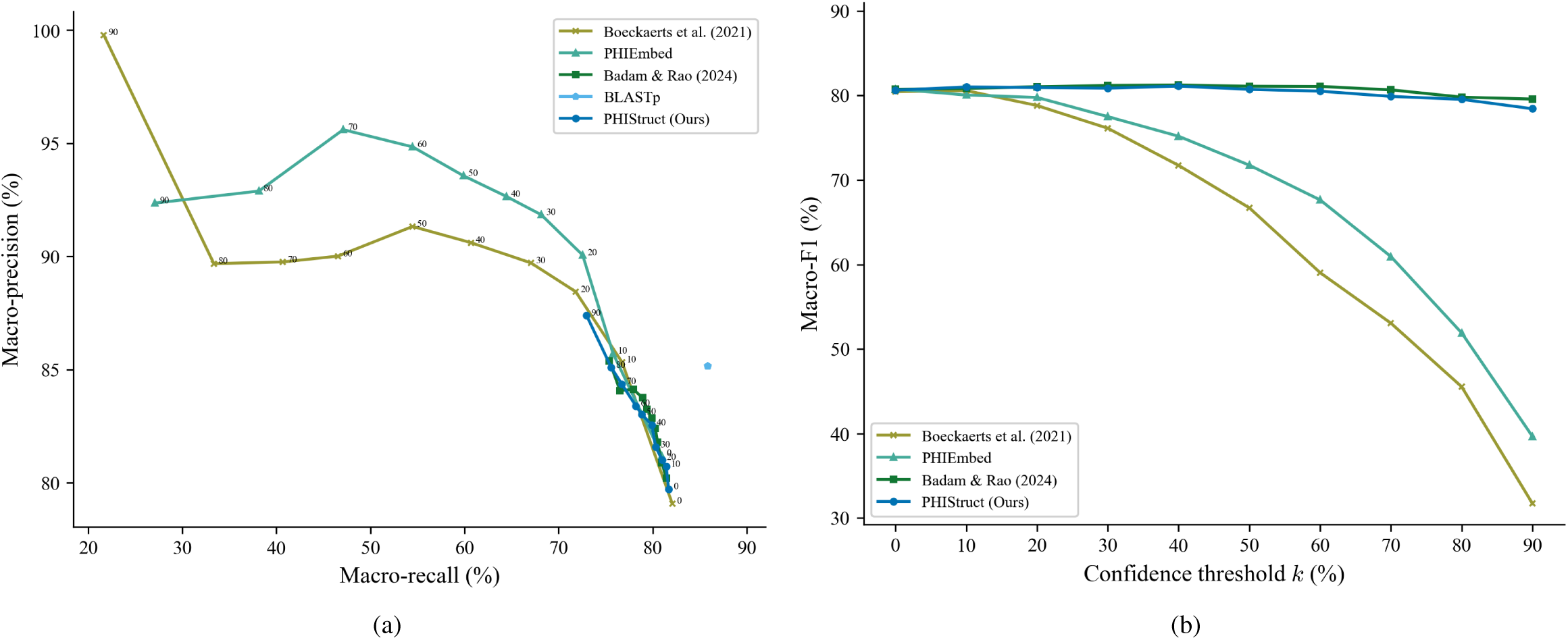
Comparison of the performance of PHIStruct with BLASTp and state-of-the-art machine learning tools that map receptor-binding proteins to host bacteria. The maximum train-versus-test sequence similarity is set to *s* = 100%. (a) Precision-recall curves. The label of each point denotes the confidence threshold *k* (%) at which the performance was measured. (b) F1 scores. Higher values of *k* prioritize precision over recall, whereas lower values prioritize recall.

**Figure A5:**
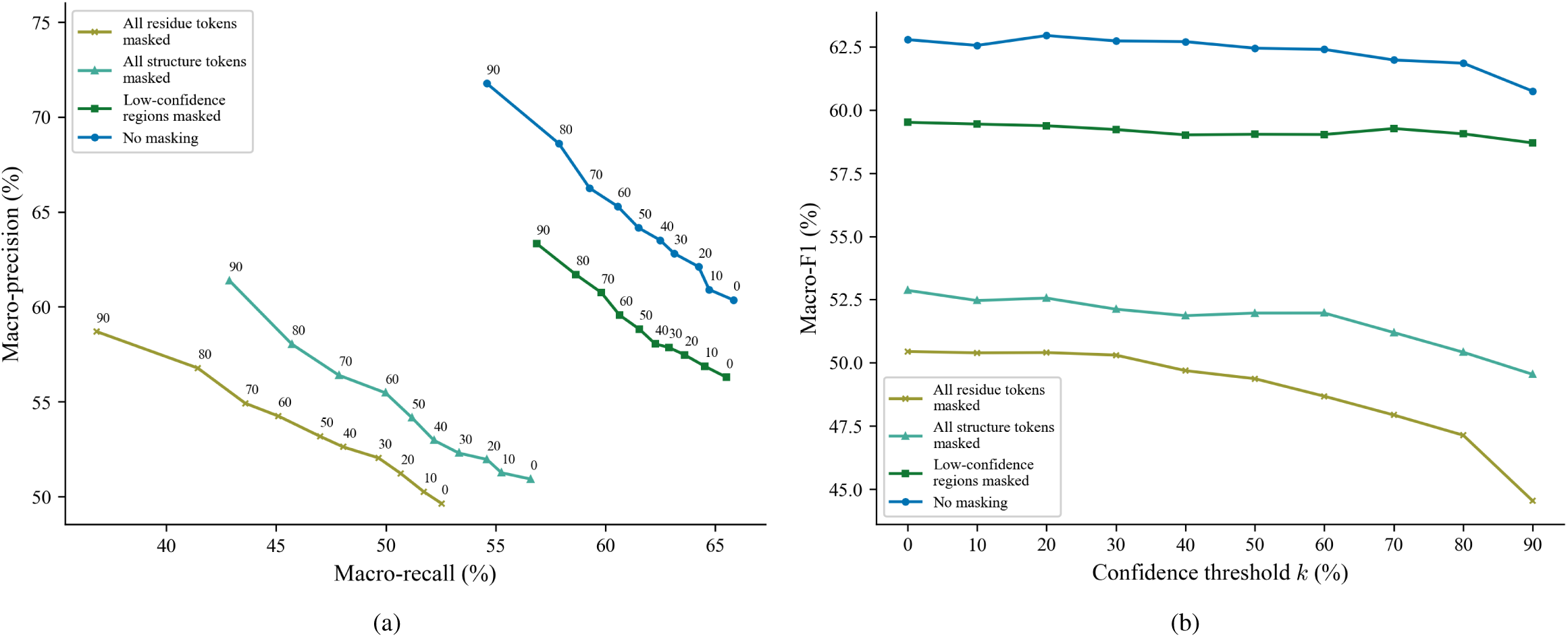
Comparison of the performance of different masking strategies for inputting proteins to SaProt. The maximum train-versus-test sequence similarity is set to *s* = 60%. (a) Precision-recall curves. The label of each point denotes the confidence threshold *k* (%) at which the performance was measured. (b) F1 scores. Higher values of *k* prioritize precision over recall, whereas lower values prioritize recall.

**Figure A6:**
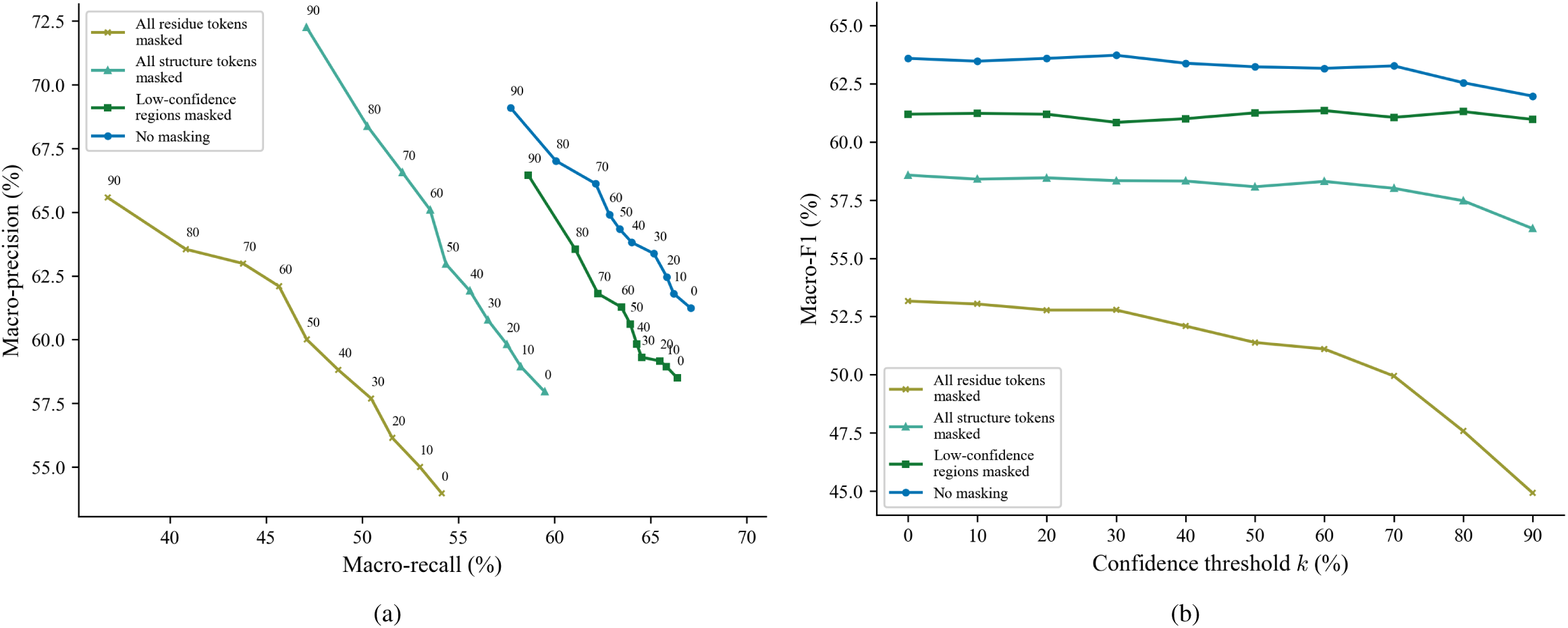
Comparison of the performance of different masking strategies for inputting proteins to SaProt. The maximum train-versus-test sequence similarity is set to *s* = 80%. (a) Precision-recall curves. The label of each point denotes the confidence threshold *k* (%) at which the performance was measured. (b) F1 scores. Higher values of *k* prioritize precision over recall, whereas lower values prioritize recall.

**Figure A7:**
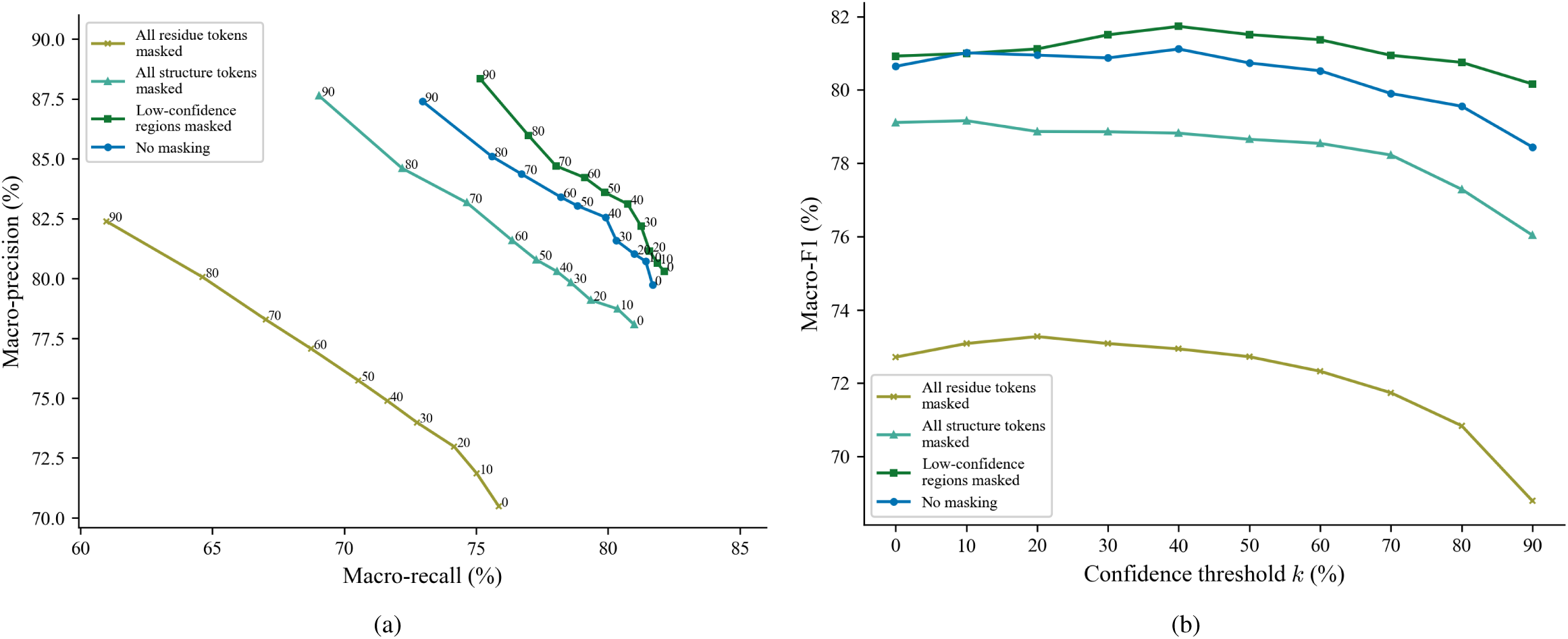
Comparison of the performance of different masking strategies for inputting proteins to SaProt. The maximum train-versus-test sequence similarity is set to *s* = 100%. (a) Precision-recall curves. The label of each point denotes the confidence threshold *k* (%) at which the performance was measured. (b) F1 scores. Higher values of *k* prioritize precision over recall, whereas lower values prioritize recall.

**Figure A8:**
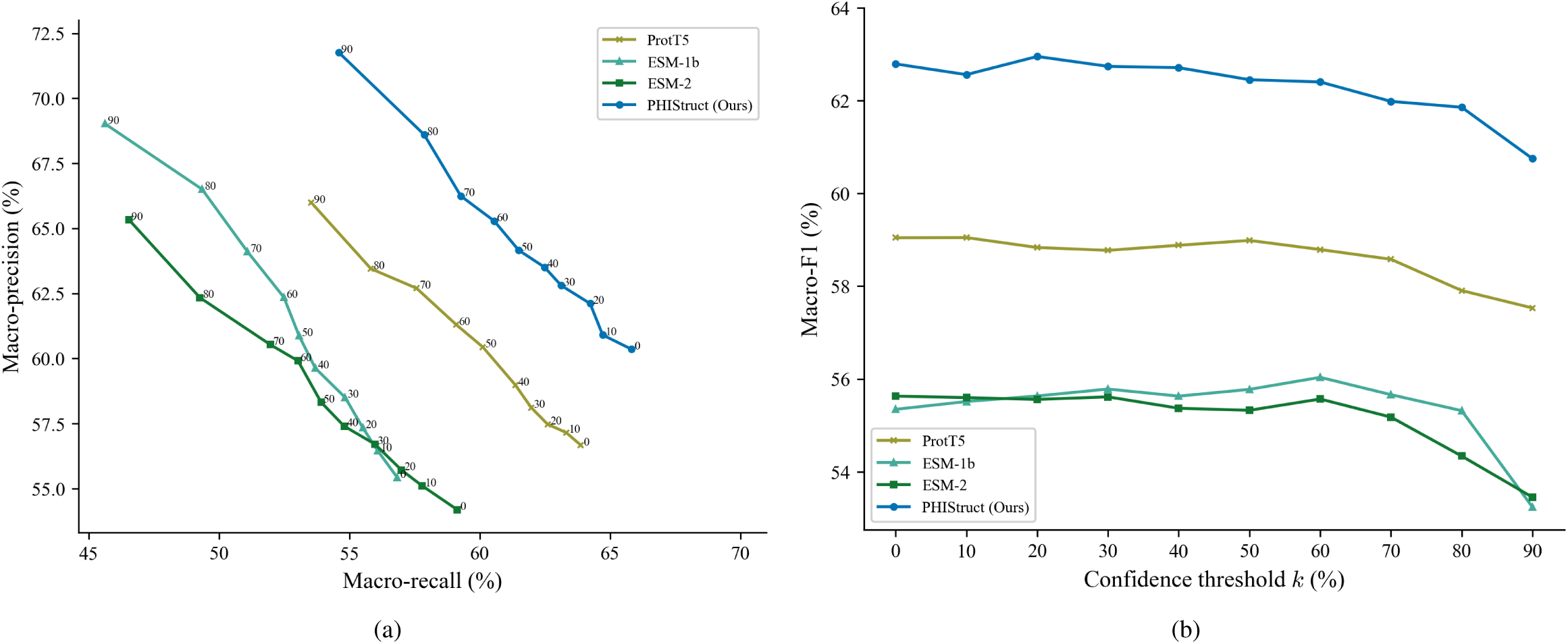
Comparison of the performance of PHIStruct with same-architecture multilayer perceptron models that take in sequence-only embeddings. The maximum train-versus-test sequence similarity is set to *s* = 60%. (a) Precision-recall curves. The label of each point denotes the confidence threshold *k* (%) at which the performance was measured. (b) F1 scores. Higher values of *k* prioritize precision over recall, whereas lower values prioritize recall.

**Figure A9:**
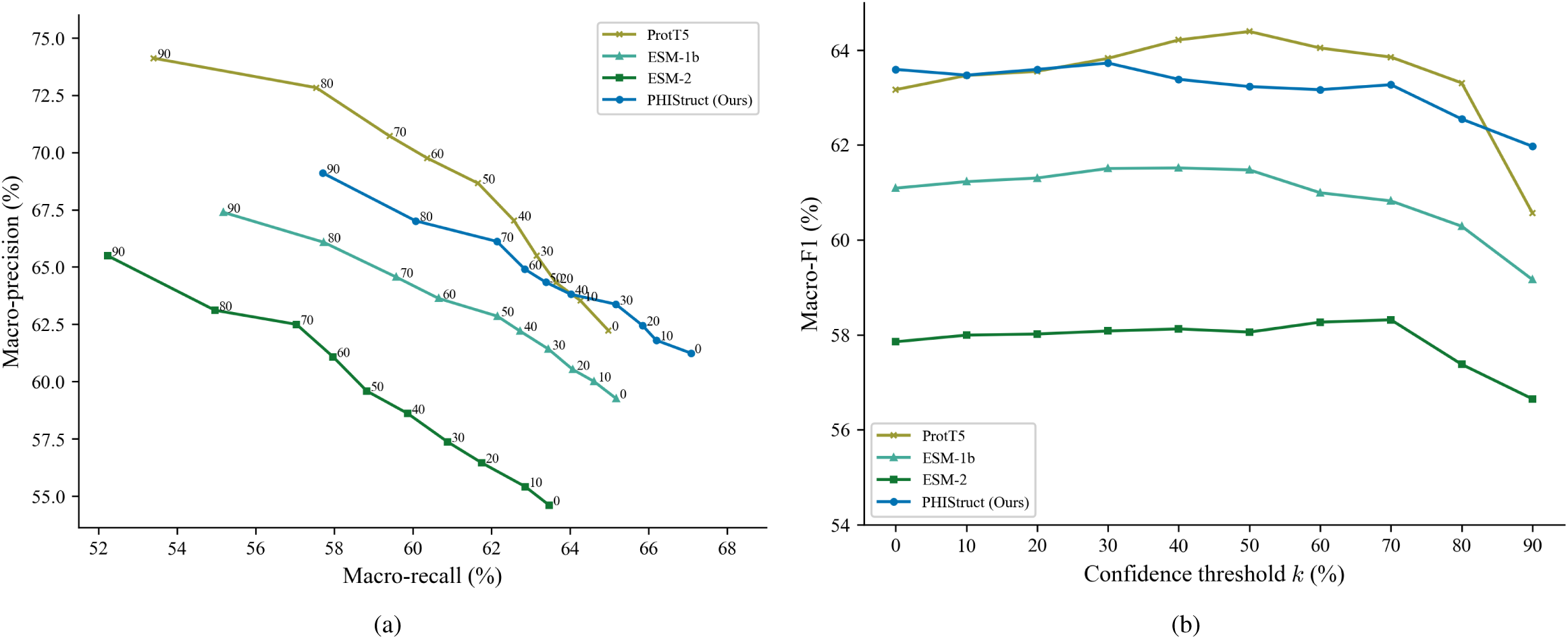
Comparison of the performance of PHIStruct with same-architecture multilayer perceptron models that take in sequence-only embeddings. The maximum train-versus-test sequence similarity is set to *s* = 80%. (a) Precision-recall curves. The label of each point denotes the confidence threshold *k* (%) at which the performance was measured. (b) F1 scores. Higher values of *k* prioritize precision over recall, whereas lower values prioritize recall.

**Figure A10:**
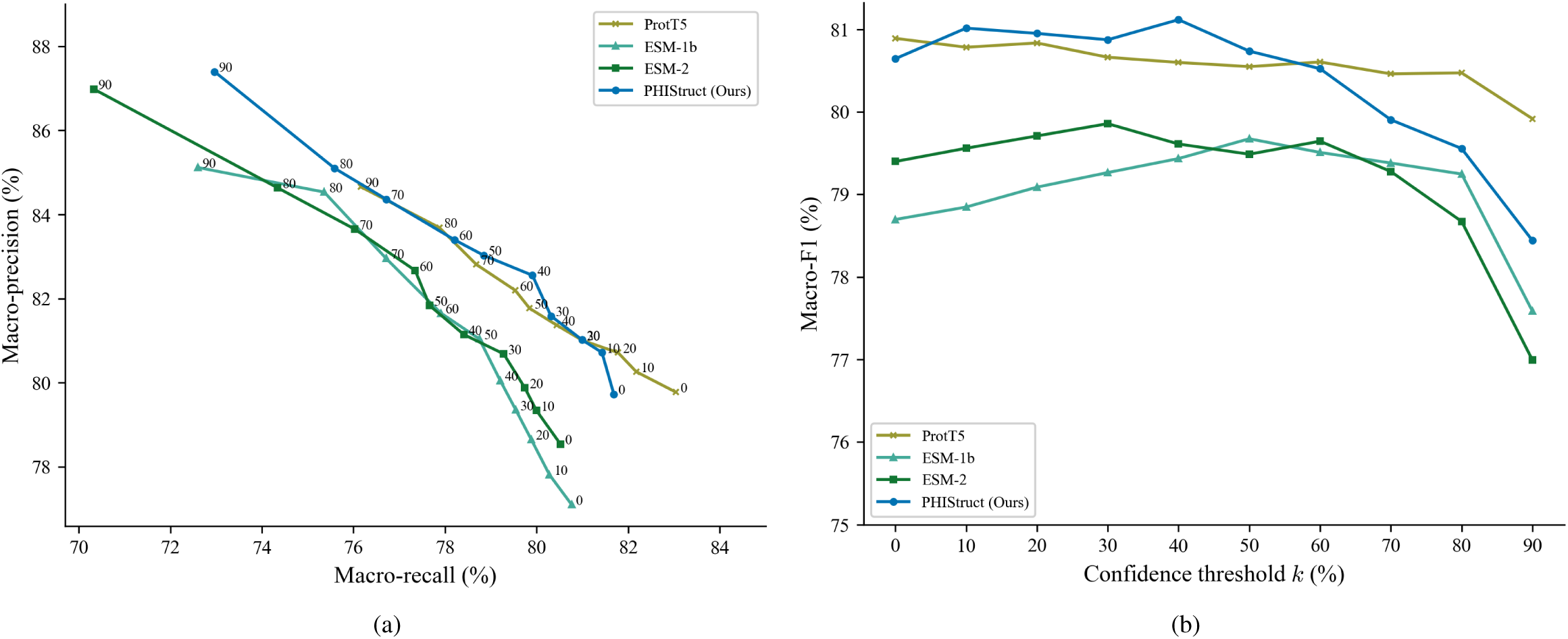
Comparison of the performance of PHIStruct with same-architecture multilayer perceptron models that take in sequence-only embeddings. The maximum train-versus-test sequence similarity is set to *s* = 100%. (a) Precision-recall curves. The label of each point denotes the confidence threshold *k* (%) at which the performance was measured. (b) F1 scores. Higher values of *k* prioritize precision over recall, whereas lower values prioritize recall.

**Figure A11:**
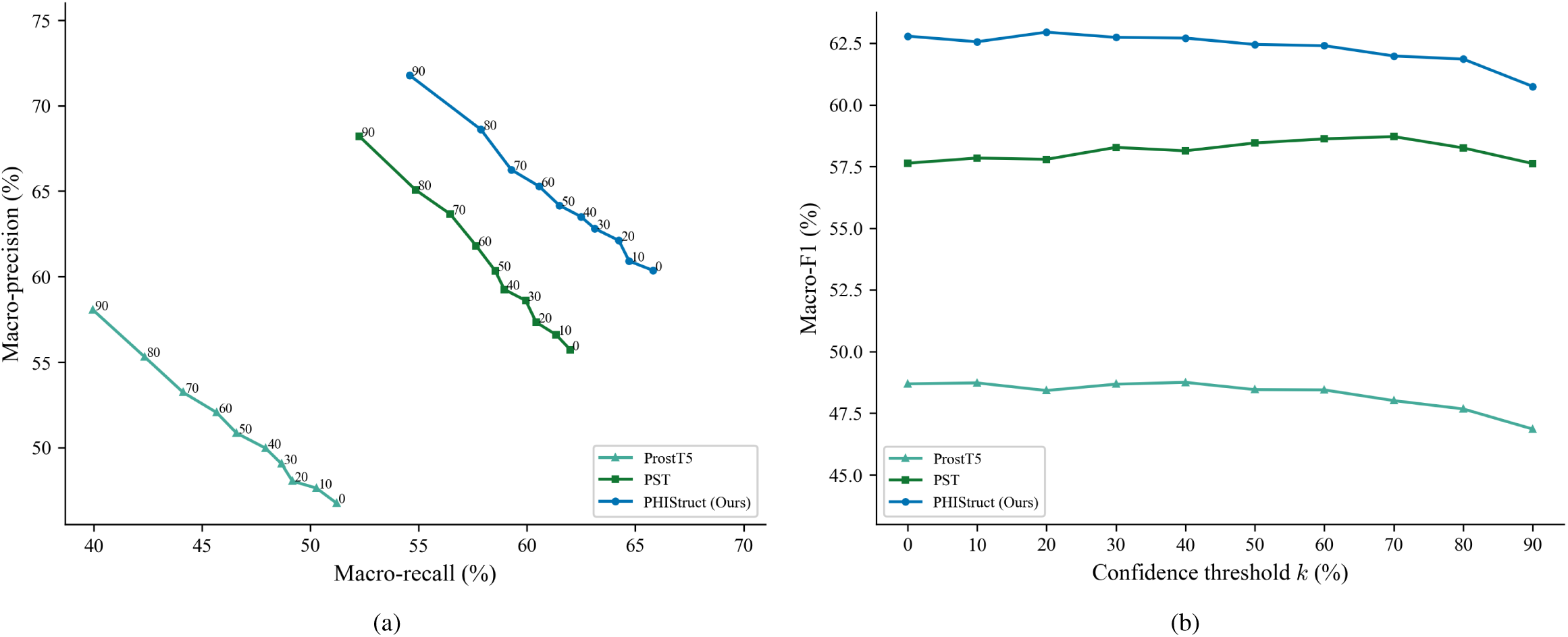
Comparison of the performance of PHIStruct with same-architecture multilayer perceptron models that take in structure-aware protein embeddings other than SaProt. The maximum train-versus-test sequence similarity is set to *s* = 60%. (a) Precision-recall curves. The label of each point denotes the confidence threshold *k* (%) at which the performance was measured. (b) F1 scores. Higher values of *k* prioritize precision over recall, whereas lower values prioritize recall.

**Figure A12:**
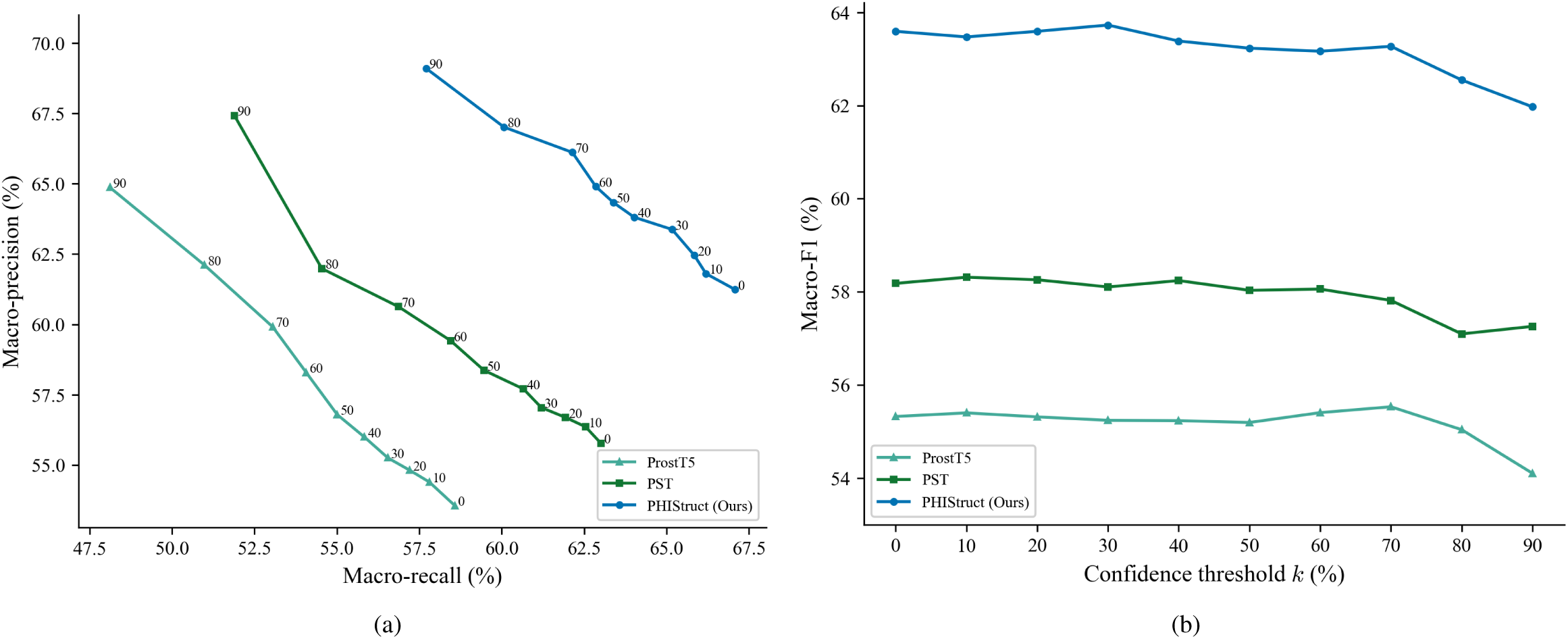
Comparison of the performance of PHIStruct with same-architecture multilayer perceptron models that take in structure-aware protein embeddings other than SaProt. The maximum train-versus-test sequence similarity is set to *s* = 80%. (a) Precision-recall curves. The label of each point denotes the confidence threshold *k* (%) at which the performance was measured. (b) F1 scores. Higher values of *k* prioritize precision over recall, whereas lower values prioritize recall.

**Figure A13:**
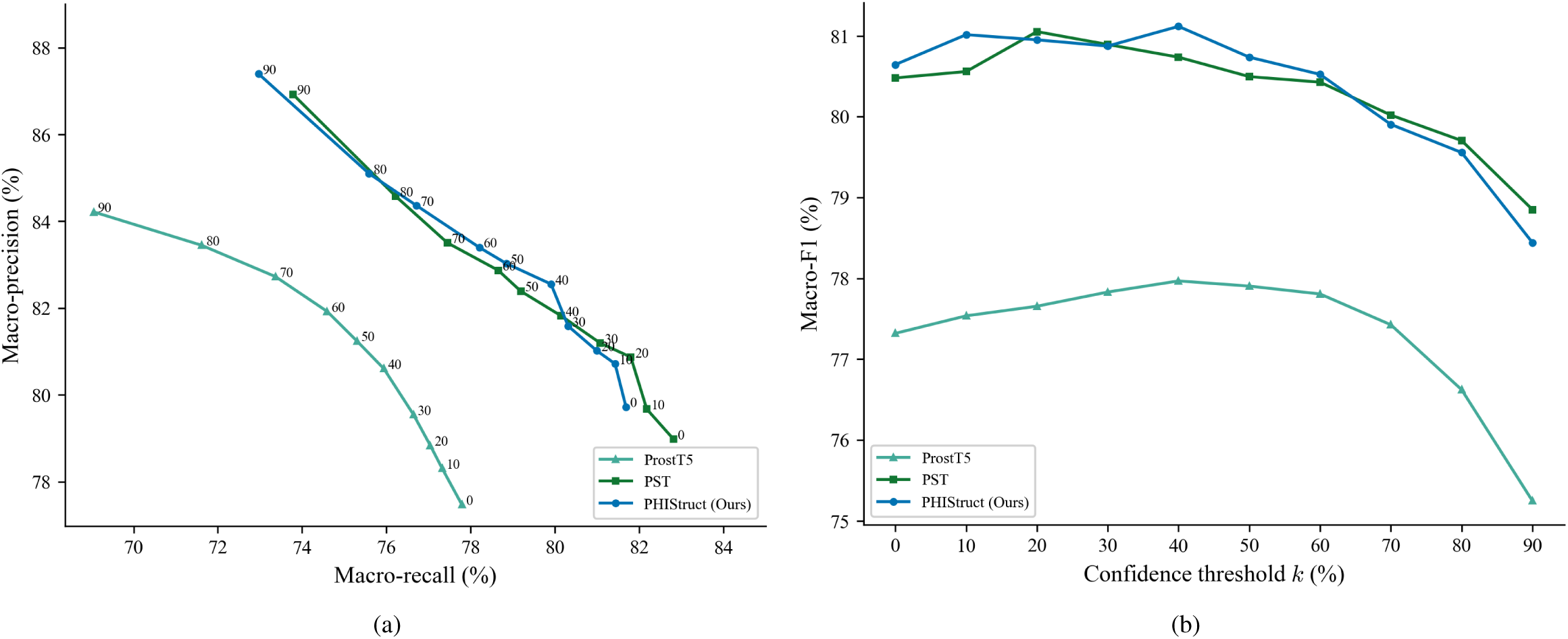
Comparison of the performance of PHIStruct with same-architecture multilayer perceptron models that take in structure-aware protein embeddings other than SaProt. The maximum train-versus-test sequence similarity is set to *s* = 100%. (a) Precision-recall curves. The label of each point denotes the confidence threshold *k* (%) at which the performance was measured. (b) F1 scores. Higher values of *k* prioritize precision over recall, whereas lower values prioritize recall.

---

**Table A1:** ColabFold parameters for protein structure prediction.

| Parameter | Value |
| --- | --- |
| Multiple sequence alignment mode | MMseqs2 (UniRef + Environmental) |
| Number of models | 1 |
| Number of recycles | 3 |
| Use of templates | Yes |
| *Amber relaxation parameter* |  |
| Maximum number of iterations | 200 |
| Maximum number of outer iterations | 3 |
| Tolerance | 2.39 |
| Stiffness | 10.0 |

**Table A2:** Per-class training and test set statistics.

| Sequence similarity $s$ | Genus | Training set | | Test set |
| --- | --- | --- | --- | --- |
|  |  | Before SMOTE-Tomek | After SMOTE-Tomek |  |
| 100% | *Enterococcus* | 119 | 2,424 | 51 |
|  | *Staphylococcus* | 365 | 2,424 | 156 |
|  | *Klebsiella* | 1,074 | 2,417 | 461 |
|  | *Acinetobacter* | 258 | 2,423 | 111 |
|  | *Pseudomonas* | 826 | 2,422 | 354 |
|  | *Enterobacter* | 272 | 2,420 | 116 |
|  | *Escherichia* | 2,424 | 2,412 | 1,040 |
|  | Others | 0 | 0 | 51 |
| 80% | *Enterococcus* | 110 | 2,120 | 60 |
|  | *Staphylococcus* | 349 | 2,120 | 172 |
|  | *Klebsiella* | 976 | 2,112 | 559 |
|  | *Acinetobacter* | 260 | 2,120 | 109 |
|  | *Pseudomonas* | 829 | 2,119 | 351 |
|  | *Enterobacter* | 221 | 2,119 | 167 |
|  | *Escherichia* | 2,120 | 2,112 | 1,344 |
|  | Others | 0 | 0 | 60 |
| 60% | *Enterococcus* | 121 | 1,875 | 49 |
|  | *Staphylococcus* | 364 | 1,875 | 157 |
|  | *Klebsiella* | 947 | 1,866 | 588 |
|  | *Acinetobacter* | 248 | 1,875 | 121 |
|  | *Pseudomonas* | 776 | 1,874 | 404 |
|  | *Enterobacter* | 192 | 1,870 | 196 |
|  | *Escherichia* | 1,875 | 1,864 | 1,589 |
|  | Others | 0 | 0 | 49 |
| 40% | *Enterococcus* | 129 | 1,668 | 41 |
|  | *Staphylococcus* | 339 | 1,668 | 182 |
|  | *Klebsiella* | 1,008 | 1,663 | 527 |
|  | *Acinetobacter* | 280 | 1,668 | 89 |
|  | *Pseudomonas* | 869 | 1,668 | 311 |
|  | *Enterobacter* | 172 | 1,666 | 216 |
|  | *Escherichia* | 1,668 | 1,661 | 1,796 |
|  | Others | 0 | 0 | 41 |

**Table A3:**
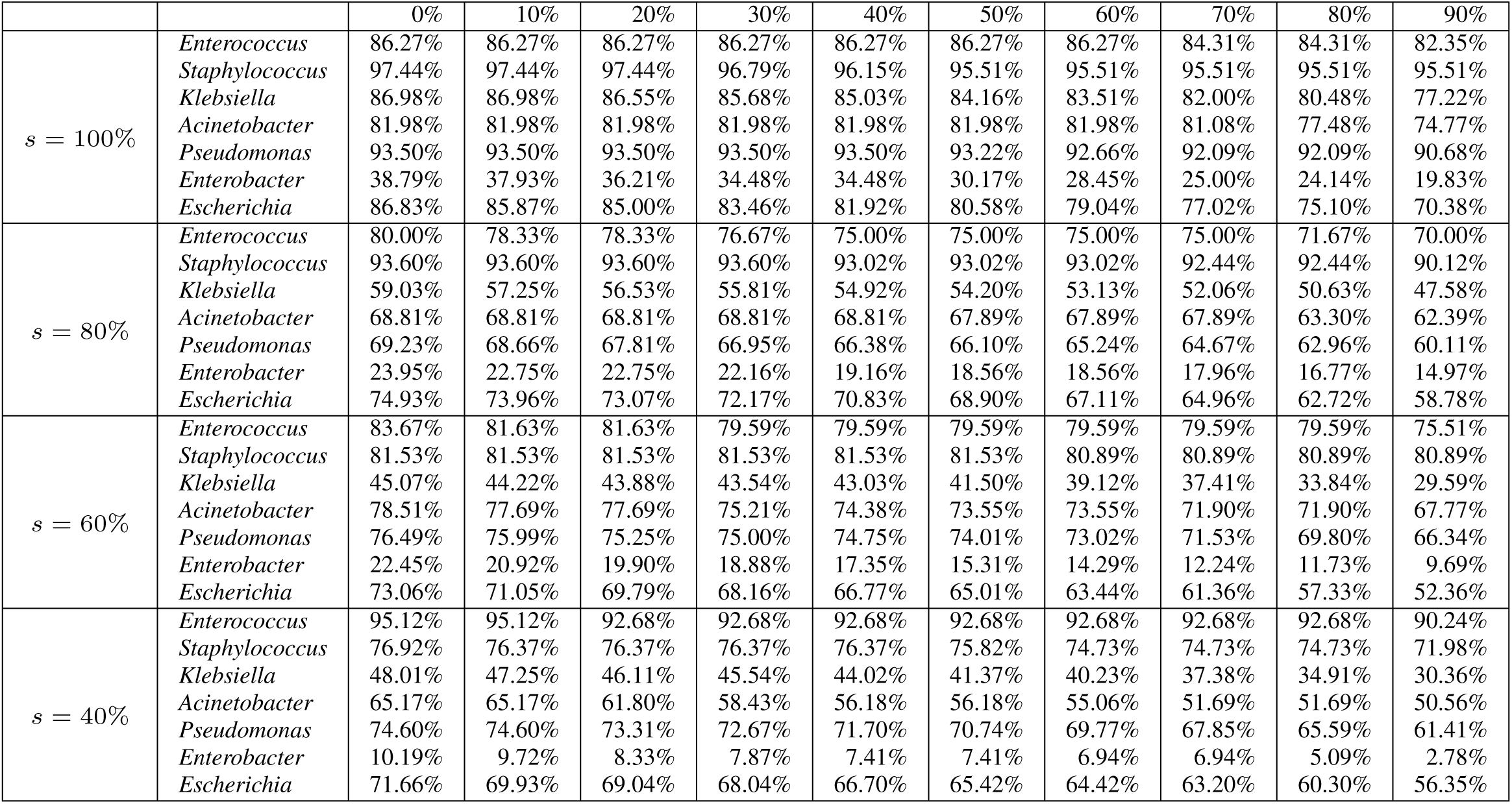
Per-class recall of PHIStruct. Lower values of the maximum train-versus-test sequence *s* indicate that the sequences in the training set are more dissimilar to those in the test set. Lower values of the confidence threshold *k* (header row) prioritize recall over precision.

**Table A4:**
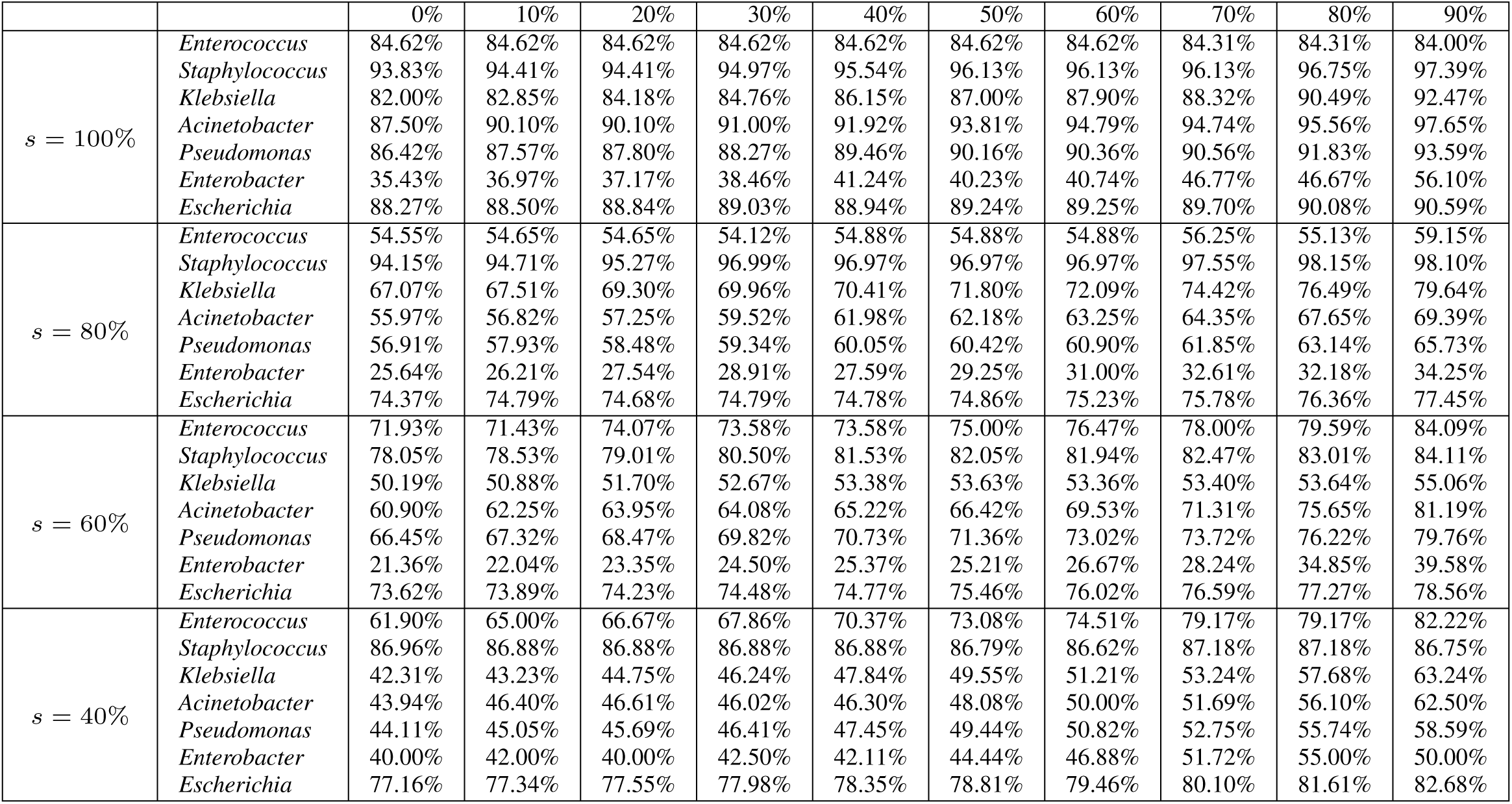
Per-class precision of PHIStruct. Lower values of the maximum train-versus-test sequence *s* indicate that the sequences in the training set are more dissimilar to those in the test set. Higher values of the confidence threshold *k* (header row) prioritize precision over recall.

**Table A5:**
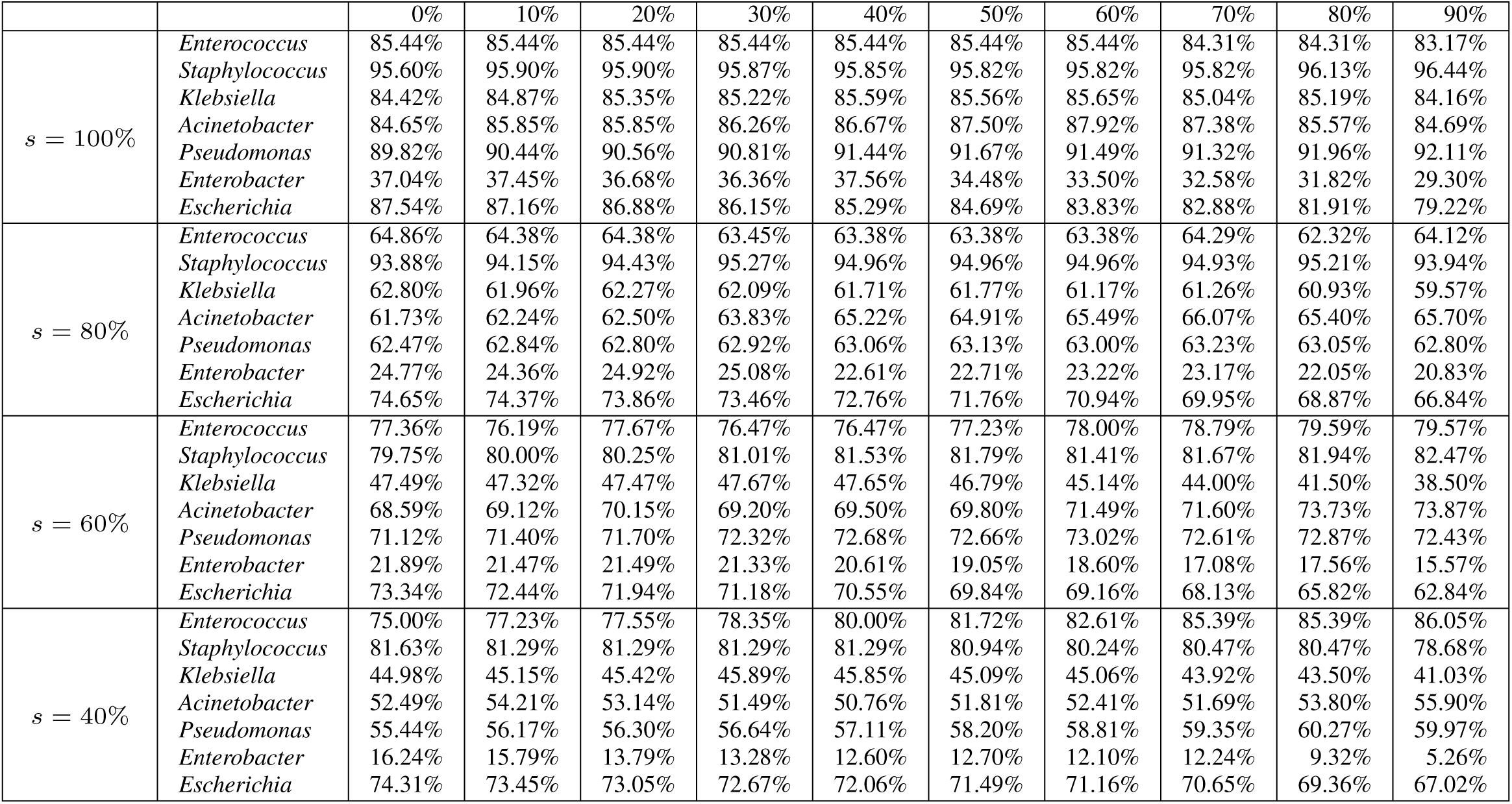
Per-class F1 of PHIStruct. Lower values of the maximum train-versus-test sequence *s* indicate that the sequences in the training set are more dissimilar to those in the test set. Higher values of the confidence threshold *k* (header row) prioritize precision over recall, whereas lower values prioritize recall.

**Table A6:** Results of the Mann-Whitney U test for investigating the structure similarity (in terms of root mean square deviation) of RBPs from phages infecting the same host versus those infecting a different host genus. For every genus, we constructed two groups, each with 500 randomly sampled RBP pairs. In the first group, both RBPs in each pair target the same genus of interest. In the second group, one of the RBPs in each pair targets the genus of interest, while the other RBP targets a different genus.

| Genus | $p$-value |
| --- | --- |
| *Enterococcus* | $4.21 \times 10^{-2}$ |
| *Staphylococcus* | $5.23 \times 10^{-7}$ |
| *Klebsiella* | $9.80 \times 10^{-2}$ |
| *Acinetobacter* | $4.99 \times 10^{-3}$ |
| *Pseudomonas* | $1.88 \times 10^{-3}$ |
| *Enterobacter* | $3.97 \times 10^{-3}$ |
| *Escherichia* | $1.59 \times 10^{-2}$ |

**Table A7:** Results of the Mann-Whitney U test for investigating the similarity (in terms of cosine distance) of the embeddings of RBPs from phages infecting the same host versus those infecting a different host genus. For every genus, we constructed two groups, each with 500 randomly sampled RBP pairs. In the first group, both RBPs in each pair target the same genus of interest. In the second group, one of the RBPs in each pair targets the genus of interest, while the other RBP targets a different genus.

| Genus | $p$-value |
| --- | --- |
| *Enterococcus* | $2.79 \times 10^{-1}$ |
| *Staphylococcus* | $1.41 \times 10^{-36}$ |
| *Klebsiella* | $1.49 \times 10^{-4}$ |
| *Acinetobacter* | $2.76 \times 10^{-6}$ |
| *Pseudomonas* | $8.36 \times 10^{-1}$ |
| *Enterobacter* | $5.62 \times 10^{-3}$ |
| *Escherichia* | $6.62 \times 10^{-4}$ |

**Table A8:** Protein language models used in benchmarking PHIStruct’s performance.

| Model | Architecture | Number of parameters | Pretraining dataset |
| --- | --- | --- | --- |
| *Sequence-only protein language models* |  |  |  |
| ProtT5 [21] | Transformer | 3 billion (ProtT5-XL) | BFD [50, 52],<br>fine-tuned on UniRef-50 [55] |
| ESM-1b [47] | Transformer | 650 million | UniRef-50 [55] |
| ESM-2 [40] | Transformer | 650 million | UniRef-50 [55] |
| *Structure-aware protein language models* |  |  |  |
| ProstT5 [28] | Transformer (based on the 3-billion-parameter version of ProtT5) | 3 billion | AlphaFold DB [61] |
| PST [16] | Graph isomorphism network and transformer (based on the 650-million-parameter version of ESM-2) | 1 billion | AlphaFold Swiss-Prot [61] |

## Notes

https://github.com/bioinfodlsu/phistruct

### Competing Interest Statement

The authors have declared no competing interest.

